# Molecular basis of one-step methyl anthranilate biosynthesis in grapes, sweet orange, and maize

**DOI:** 10.1101/2024.06.10.598330

**Authors:** Michael A. Fallon, Hisham Tadfie, Aracely P. Watson, Madeline M. Dyke, Christopher Flores, Nathan Cook, Zhangjun Fei, Cynthia K. Holland

## Abstract

Plants synthesize an array of volatile compounds, many of which serve ecological roles in attracting pollinators, deterring herbivores, and communicating with their surroundings. Methyl anthranilate is an anti-herbivory defensive volatile responsible for grape aroma that is emitted by several agriculturally relevant plants, including citrus, grapes, and maize. Unlike maize, which uses a one-step anthranilate methyltransferase, grapes have been thought to use a two-step pathway for methyl anthranilate biosynthesis. By mining available transcriptomics data, we identified two anthranilate methyltransferases in *Vitis vinifera* (wine grape), as well as one ortholog in ‘Concord’ grape. Many angiosperms methylate the plant hormone salicylic acid to produce methyl salicylate, which acts as a plant-to-plant communication molecule. Because the *Citrus sinensis* (sweet orange) salicylic acid methyltransferase can methylate both anthranilate and salicylic acid, we used this enzyme to examine the molecular basis of anthranilate activity by introducing rational mutations, which identified several active site residues that increase activity with anthranilate. Reversing this approach, we introduced mutations that imparted activity with salicylic acid in the maize anthranilate methyltransferase, which uncovered different active site residues from those in the citrus enzyme. Sequence and phylogenetic analysis revealed that one of the *Vitis* anthranilate methyltransferases shares an ancestor with jasmonic acid methyltransferases, similar to the anthranilate methyltransferase from strawberry (*Frageria* sp.). Collectively, these data demonstrate the molecular mechanisms underpinning anthranilate activity across methyltransferases and identified one-step enzymes by which grapes synthesize methyl anthranilate.

**Significance Statement:** While the two-step pathway responsible for the biosynthesis of the grape aroma molecule, methyl anthranilate, has remained incomplete in *Vitis* spp., we identified two one-step anthranilate methyltransferases in wine and one in ‘Concord’ grapes that can methylate the tryptophan pathway intermediate anthranilate. Tracing the molecular basis of anthranilate activity in the maize and sweet orange methyltransferases uncovered distinct active site amino acids that impart substrate specificity.

## INTRODUCTION

Plants synthesize an array of volatiles that impart distinct aromas (Dudareva *et al*., 2004, Dudareva *et al*., 2006). These volatiles serve biological and ecological roles that enable plants to attract pollinators and seed dispersers, repel herbivores, and communicate with their surroundings, including other plants (Schuman, 2023). Plant volatiles also have economic value as aromatic and flavoring agents in the food, beverage, cosmetic, fragrance, and pharmaceutical industries (Baldwin, 2010). Many flowering plants synthesize the wintergreen aroma compound methyl salicylate (MeSA), which is the methyl ester of the plant hormone salicylic acid (SA) (Figure 1) (Effmert *et al*., 2005). Methyl anthranilate (MeAA) is the methyl ester of the tryptophan pathway intermediate anthranilate (AA) and is responsible for the recognizable aroma of grape (Wang and De Luca, 2005, Li *et al*., 2023). In addition to being volatiles, methylating plant hormones, such as SA, jasmonate, and indole 3-acetic acid (auxin), inactivates them and modulates their activities (Westfall *et al*., 2013).

**Figure 1.**
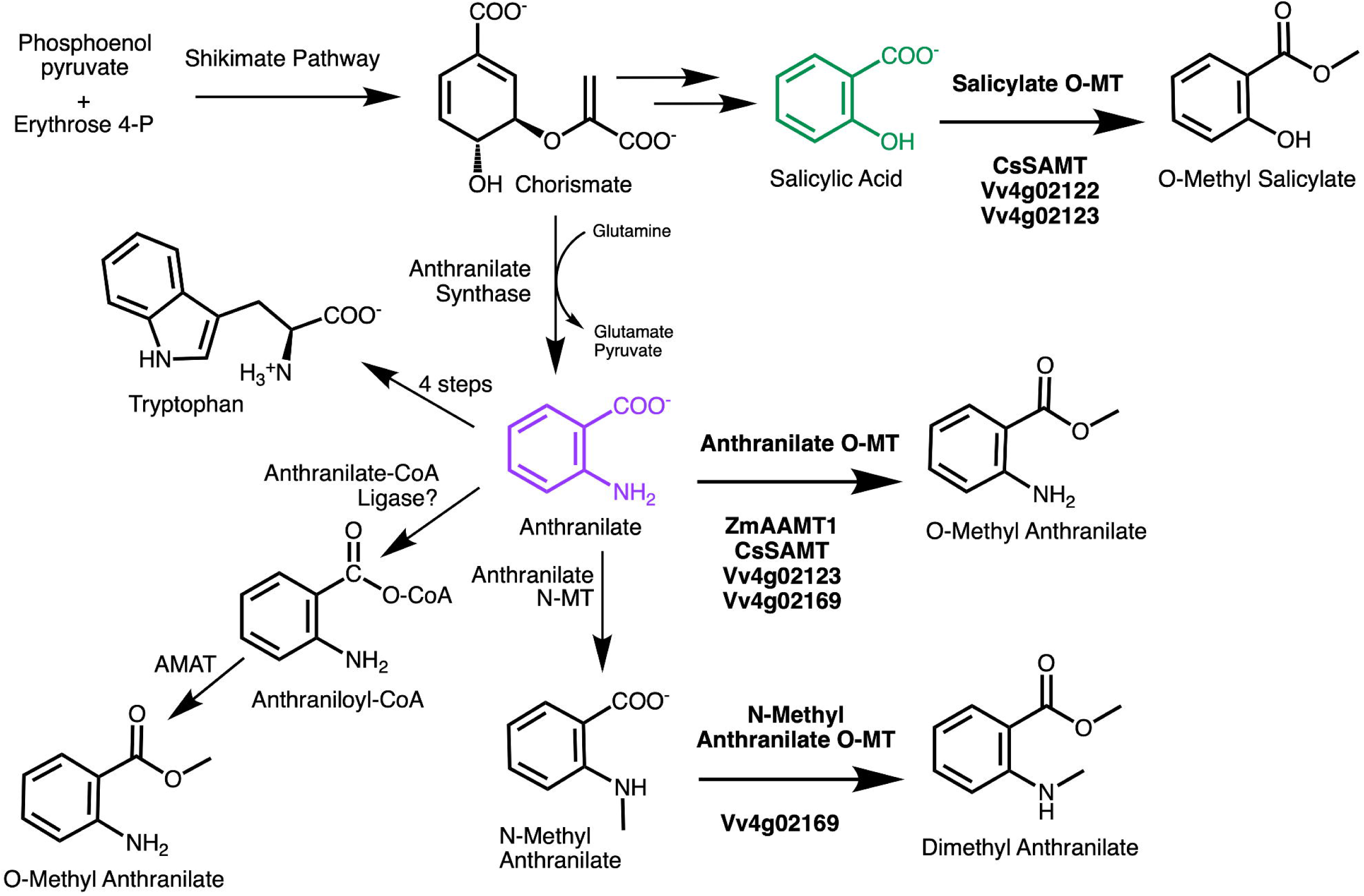
Schematic diagram of the biosynthesis and fates of anthranilate in plants. The names of enzymes investigated here in citrus (Cs), grapes (Vv), and maize (Zm) are displayed in bold below the arrow.

MeAA-producing species include many agriculturally relevant crops, such as maize (*Zea mays)*, alfalfa (*Medicago truncatula*), fox grapes (*Vitis labrusca*), strawberries (*Fragaria* spp.), citrus (*Citrus* spp.), and soybeans (*Glycine max*) (Wang and De Luca, 2005, Kollner *et al*., 2010, Lin *et al*., 2013, Pillet *et al*., 2017, Gonzalez-Mas *et al*., 2019, Pollier *et al*., 2019). MeSA and MeAA play important roles in plant defense against herbivores. When plants are under attack by aphids, they emit volatile MeSA, which can travel long distances to neighboring plants that perceive MeSA, demethylate it, and use the resulting SA to regulate transcriptional responses and elicit defenses (Gong *et al*., 2023). Citrus plants that are infected with the psyllid-vectored bacterial pathogen *Candidatus* Liberibacter asiaticus (CLas), the causal agent of citrus greening (huanglongbing), release high levels of MeSA, while uninfected plants release high levels of MeAA (Mann *et al*., 2012). MeSA, but not MeAA, attracts the psyllid (*Diaphorina citri*) to the plant, which in turn may promote the spread of the pathogen. Strawberries that are resistant to *Drosophila suzukii* pest flies produce high concentrations of MeAA, which attracts the egg-laying adults but then leads to reduced hatching rates of their eggs (Bracker *et al*., 2020). Maize synthesizes and releases a volatile blend that contains MeSA and MeAA in response to insect damage, which then attracts wasps that parasitize the insect herbivores (Turlings *et al*., 1990, von Merey *et al*., 2013). Exogenous application of MeAA on crops has also been shown to be effective in attracting natural enemies of the herbivores to protect plants from further damage (Simpson *et al*., 2011). Commercially, MeAA is also an effective bird repellant, and MeAA sprays are available for deterring birds from orchards and turf (Mason *et al*., 1989, Avery *et al*., 1995, Mikiciuk *et al*., 2021).

Although humans frequently associate the aroma of MeAA with grapes, the biosynthetic pathway has remained incomplete in *Vitis* spp. (Wang and De Luca, 2005). While MeAA is high in ‘Concord’ grape juices, this compound contributes to a ‘foxy’ aroma that is undesirable in wine and is found at low or undetectable levels in wine grapes (*Vitis vinifera*) (Sun *et al*., 2011, Lin *et al*., 2019). In ‘Washington Concord’ grape, an anthraniloyl-coenzyme A (CoA): methanol acyltransferase (AMAT) that condenses anthraniloyl-CoA and methanol into MeAA has been identified (Figure 1) (Wang and De Luca, 2005). However, the two-step MeAA pathway remains incomplete as the first pathway enzyme—anthranilate-CoA ligase—remains to be discovered.

Maize and strawberry instead use a one-step S-adenosyl-L-methionine (SAM)-dependent methyltransferase for MeAA biosynthesis (Kollner *et al*., 2010, Pillet *et al*., 2017), and in citrus, a one-step SA methyltransferase (SAMT) that can methylate anthranilate has been identified (Huang *et al*., 2016) (Figure 1). *SAMT* orthologs are found in most angiosperm orders (Dubs *et al*., 2022), and while MeAA has been detected in the headspace floral volatiles of at least nineteen plant families (Knudsen *et al*., 2006), AA-using enzymes remain largely unidentified in MeAA-producing plants. Because SA and AA are structurally similar and differ only in that the *ortho* hydroxyl on SA is an amine on AA, we hypothesized that active site residues in SA- and/or AA-using methyltransferase would confer activity with and specificity for one or both substrates. We first sought to identify and functionally characterize AA methyltransferases (AAMTs) in grapes (*Vitis* spp.) by mining available metabolomics, transcriptomics, and genomic resources. Furthermore, we set out to trace the molecular mechanisms of AA recognition in SABATH (<u>s</u>alicylic <u>a</u>cid, <u>b</u>enzoic <u>a</u>cid, <u>th</u>eobromine) methyltransferases in citrus and maize.

## RESULTS AND DISCUSSION

### Identification of one-step S-adenosyl-methionine-dependent anthranilate methyltransferases in grapes

While citrus, maize, and strawberries synthesize MeAA using a one-step methyltransferase, *Vitis* spp. are thought to synthesize MeAA via a two-step two-enzyme pathway, and only the second enzyme in this pathway, an AMAT, has been characterized (Wang and De Luca, 2005) (Figure 1). AMAT had a turnover rate of 1.32 min^-1^ and displayed 26-fold higher relative activity with benzyl alcohol compared to methanol, suggesting that additional enzymes may be responsible for MeAA synthesis in grape (Wang and De Luca, 2005). Although the *AMAT* gene expression profile corresponds to MeAA levels in berries during ripening (Yang *et al*., 2020), we hypothesized that grapes may also have a SABATH methyltransferase that could catalyze the methylation of AA. Published metabolomics data showed that MeAA levels increase throughout fruit ripening, and this same study used transcriptomics to identify genes that are expressed in ripening fruits in various *Vitis* wine accessions and in ‘Concord’ (Yang *et al*., 2020). From these available ‘omics’ data, we selected four candidate methyltransferases that have the highest expression in ‘Concord’ fruit, where MeAA accumulates (Data S1).

To determine whether the four candidate methyltransferases from *V. vinifera* were active with AA, each enzyme was assayed *in vitro,* and this confirmed that two of the candidates from *V. vinifera* (Vv4g02123 and Vv4g02169) methylated AA (Figure 2a). Vv4g02123, had similar activities with 1 mM AA, SA, and BA, but the Km values could not be captured because the activity was linear to 1 mM, indicating that the enzyme did not have a preference for these substrates (Figures 2b and 2d; Figure S1). Unexpectedly, when assayed with N-methyl AA (N-MeAA), Vv4g02123 had a low Km value of 12 μM, suggesting that the enzyme prefers N-MeAA as a substrate, forming dimethyl AA (DiMeAA) (Figure 2a and 2c). However, Vv4g02123 had 2-fold lower activity with 1 mM N-MeAA compared to its activity with 1 mM AA. The products of Vv4g02123— MeAA and DiMeAA—were confirmed by GC-MS in comparison to authentic standards (Figure S2 and S3).

**Figure 2.**
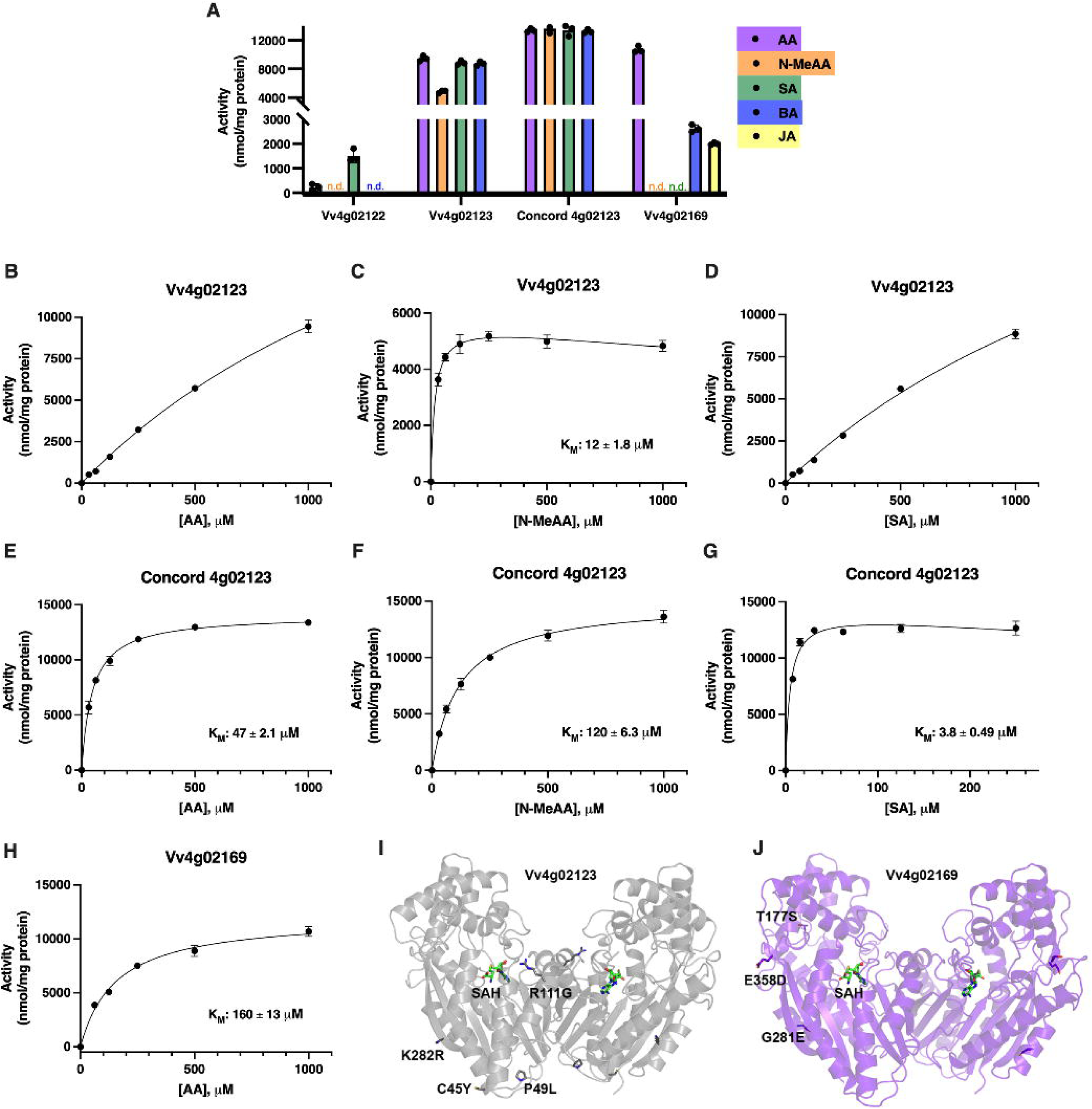
Identification of two anthranilate methyltransferases in *Vitis vinifera* and ‘Concord’. (A) Enzyme activity across *Vitis vinifera* (Vv) and ‘Concord’ methyltransferases with 1 mM substrates. Only Vv4g2169 was assayed with JA. (B) Kinetics plot of Vv4g02123 with varied anthranilate (AA), (C) varied N-methyl anthranilate (N-MeAA) fit to a substrate inhibition kinetics curve, and (D) varied salicylic acid (SA). (E) Kinetics plot of the ‘Concord’ ortholog of Vv4g02123 with varied AA, (F) varied N-MeAA, and (G) varied SA, which was fit to a substrate inhibition kinetics curve. (H) Kinetics plot of Vv4g02169 with varied AA. (I) Structural differences between Vv4g02123 and (J) Vv4g02169 in *V. vinifera* and their orthologs in ‘Concord.’ SAH (green) is positioned in the active site of each dimer. For all points, n=3; where error bars are not visible, standard deviation is too small to visualize.

DiMeAA is also a volatile, but very little is known about its biosynthesis in plants. Synthesizing DiMeAA would require a pool of N-MeAA to use as a substrate, which may be synthesized by an anthranilate *N-*methyltransferase (Figure 1). While an anthranilate N-methyltransferase has been identified in common rue (*Ruta graveolens*) (Rohde *et al*., 2008), one remains to be identified in *Vitis* spp. DiMeAA (also methyl *N*-methyl aminobenzoate) has been reported in floral volatiles of plants in the Orchidaceae and Rutaceae but has yet to be identified in Vitaceae species (Knudsen *et al*., 2006).

The second candidate that was active with AA, Vv4g02169, had a low Km value of 160 μM for AA, suggesting that this was the preferred substrate (Figure 2h). Also, Vv4g02169 had 5-fold higher activity with AA compared to BA, and activity was not detected with SA or N-MeAA (Figure 2a). We hypothesized that another candidate, Vv4g02122, would function as an AA methyltransferase based on high gene expression in ‘Concord’ berries (Yang *et al*., 2020), but it methylated SA, had minimal activity with AA, and was inactive with BA (Figure 2a; Figure S1). The fourth candidate, Vv12g00725, did not methylate SA, AA, BA, or N-MeAA when assayed *in vitro*. Therefore, we have identified two *V. vinifera* enzymes that produce MeAA, and one of which also produces DiMeAA.

While these enzymes were from wine grapes (*V. vinifera*), MeAA is found at low concentrations in wine grapes and at high concentrations in ‘Concord’, which is a hybrid of *Vitis labrusca* with roughly one third of its genome from *V. vinifera* (Sawler et al., 2013). Since published transcriptomics data indicates all of the identified methyltransferases are expressed in both *V. vinifera* and ‘Concord’ berries during ripening, we sought to understand the genomic variation that accounts for the differences in their MeAA profiles. To illuminate differences in gene expression and/or enzyme activity, we compared the promoter and gene sequences of the two AA-using methyltransferases, *Vv4g02123* and *Vv4g02169*, to those of *V. labrusca* and ‘Concord’. In the promoter region, there is a 166-bp deletion that is 784 bp upstream of the start codon and a 23-bp insertion that is 686 bp upstream of the start codon in *V. labrusca* and ‘Concord’ orthologs when compared to the promoter of *Vv4g02123* (Figure S4). In the promoter of the ‘Concord’ ortholog of *Vv4g02169*, there is an AT-rich 32-bp insertion 1 kb upstream of the start codon that is not found in either *V. labrusca* or *V. vinifera*, as well as other small insertions and deletions that are not found in either parent (Figure S5). These promoter differences may contribute to differences in gene expression that lead to altered MeAA biosynthesis across grape cultivars.

In addition to changes in gene expression, mutations in the coding sequence could lead to altered enzyme activity. The coding sequence of the ‘Concord’ ortholog of *Vv4g02123* is conserved in *V. labrusca* but has four single nucleotide differences that cause amino acid changes in comparison to *V. vinifera* (C45Y, P49L, R111G, and K282R) (Figure 2i and Figure S6). Cys45, Pro49, and Lys282 are positioned away from the active site, while Arg111 is located within a helix at the dimer interface. The ‘Concord’ and *V. labrusca* orthologs of *Vv4g02169* also have conserved coding sequences; however, there is an in-frame three-bp deletion in the *V. labrusca* gene that deletes Lys4 (Figure S7). The *V. vinifera* Vv4g02169 and its ‘Concord’ ortholog differ by three amino acids (T177S, G281E, and E358D) (Figure 2j) that are positioned away from the active site and the dimer interface. While these amino acid differences were outside the active site, we purified and assayed the ‘Concord’ ortholog of 4g02123 and 4g02169 to determine whether this variation would impact activity.

When we assayed the ‘Concord’ ortholog of 4g02123, it retained activity with AA and had a low Km value of 47 μM, which is greatly reduced in comparison to the *V. vinifera* ortholog (Figure 2e). While the *V. vinifera* ortholog had a very low Km value with N-MeAA, the ‘Concord’ ortholog had a 10-fold increased Km value of 120 μM with N-MeAA, suggesting that the ‘Concord’ enzyme has relaxed preference for N-MeAA (Figure 2f) relative to the *V. vinifera* enzyme (Figure 2c). The lowest Km value of any substrate with the ‘Concord’ 4g02123 was 3.8 μM with SA, followed by 23 μM with BA (Figure 2g; Figure S2). Although there are only three minor amino acid changes between 4g02169 in *V. vinifera* and ‘Concord’, the ‘Concord’ ortholog was inactive *in vitro*, and MeAA could not be detected in the media from *E. coli* expressing the gene or single mutants of the gene. This was surprising, especially when considering that these three amino acids are positioned outside of the active site and away from the dimer interface (Figure 2j). Therefore, the ‘Concord’ ortholog of 4g02123 likely represents the primary one-step methyltransferase for MeAA biosynthesis, and the 166-bp deletion within one kb of the start codon may account for differences in MeAA profiles across grape accessions (Figure S4).

### Structure-guided analysis of AA activity in the sweet orange SAMT

Previous efforts to trace the substrate specificity for AA used the maize AAMT1, which found an active site mutation that conferred activity with SA, Y246W (Kollner *et al*., 2010). Because CsSAMT can methylate SA, AA, and BA and its expression correlates with MeAA presence in citrus flowers (Azam *et al*., 2013, Huang *et al*., 2016), we reasoned that this was a good candidate for identifying additional active site residues that contribute to AA recognition and activity in a SAMT. To identify putative active site residues that may be responsible for AA activity, we generated AlphaFold structural models of the CsSAMT and overlaid them with a solved structure of the *Clarkia breweri* SAMT, which had been co-crystalized with S-adenosyl-L-homocysteine (SAH) and SA (Zubieta *et al*., 2003). Active site residues that varied between functionally characterized AAMTs and methyltransferases that use SA or BA were identified as candidates for site-directed mutagenesis (Figure 3). These comparisons led us to mutate six variable active site residues to examine their role in AA activity: Gln159, Gln263, Cys319, Met320, Ala324, and Phe359 (Table 1).

**Figure 3.**
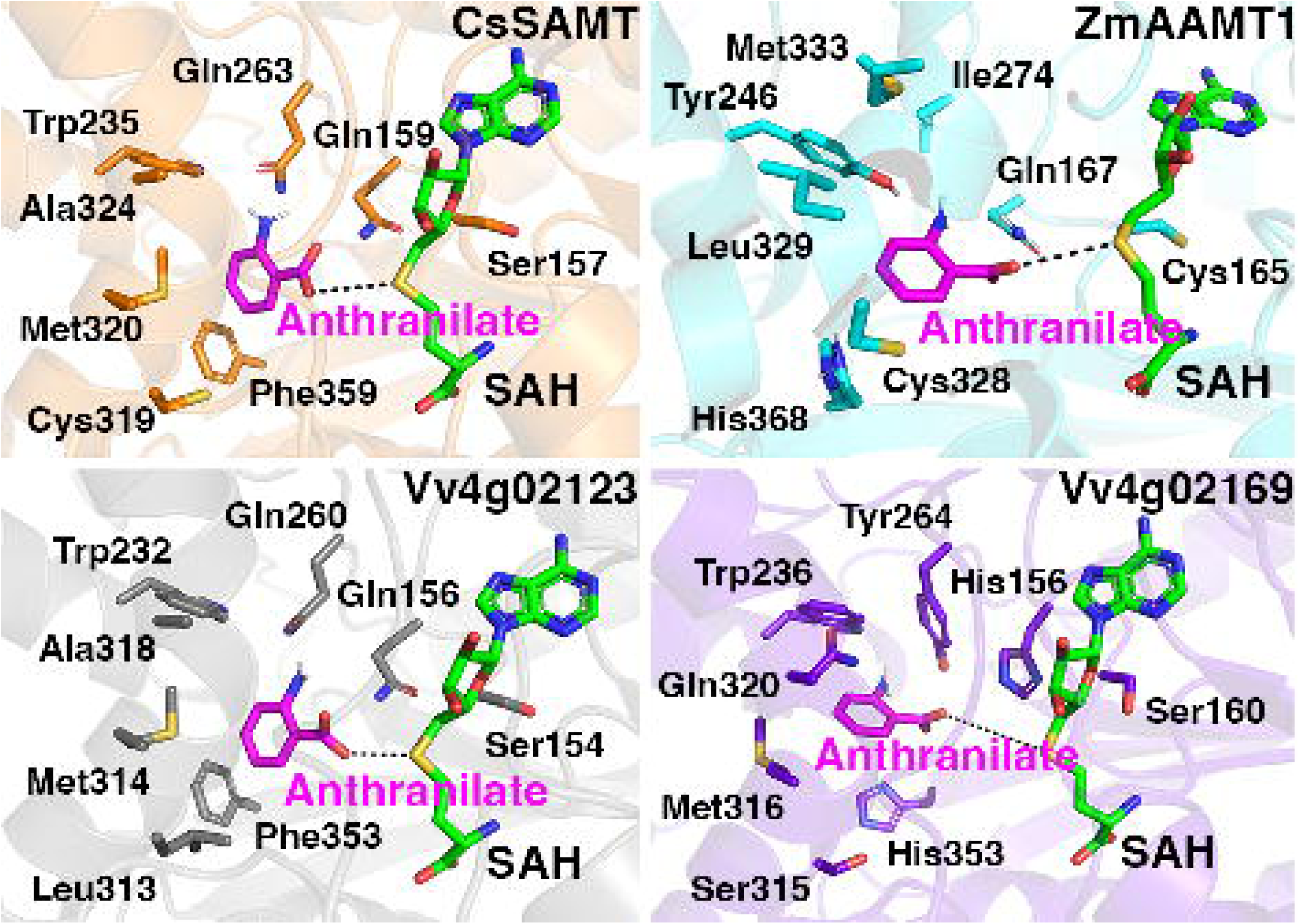
Active site comparisons of four enzymes that generate O-methyl anthranilate: *Citrus sinensis* SAMT (CsSAMT), *Zea mays* AAMT1 (ZmAAMT1), *Vitis* Vv4g02123, and *Vitis* Vv4g02169. Anthranilate (magenta) was docked into each active site, and SAH (green) was overlaid from a solved structure (PDB: 1M6E).

**Table 1.** Activities of CsSAMT wild type and single mutants with salicylic acid, anthranilate, and benzoic acid. Activities with 1 mM substrates are reported because the data were generated from endpoint assays. n= 3 replicates; S.E. = standard error; S.D. = standard deviation; n.d. = none detected.

| Protein | Salicylic Acid |  | Anthranilate |  | Benzoic Acid |  |
| --- | --- | --- | --- | --- | --- | --- |
| | Km value ± S.E. (μM) | Activity with 1 mM ± S.D. (nmol/mg) | Km value ± S.E. (μM) | Activity with 1 mM ± S.D. (nmol/mg) | Km value ± S.E. (μM) | Activity with 1 mM ± S.D. (nmol/mg) |
| CsSAMT | 105 ± 11 | 8000 ± 340 | n.d. | 870 ± 250 | n.d. | 1900 ± 280 |
| SAMT-Q159H | 390 ± 33 | 3300 ± 250 | n.d. | 160 ± 51 | n.d. | 1300 ± 35 |
| SAMT-Q263C | 840 ± 78 | 5800 ± 110 | 480 ± 75 | 1600 ± 101 | n.d. | 480 ± 170 |
| SAMT-Q263Y | 480 ± 78 | 6800 ± 900 | 230 ± 15 | 2900 ± 45 | n.d. | 150 ± 32 |
| SAMT-C319S | 180 ± 10 | 10,300 ± 130 | 790 ± 190 | 1500 ± 82 | n.d. | 760 ± 110 |
| SAMT-M320I | 200 ± 10 | 9400 ± 260 | n.d. | 1400 ± 110 | n.d. | 2600 ± 180 |
| SAMT-M320L | 560 ± 42 | 5200 ± 140 | n.d. | 1200 ± 23 | n.d. | 310 ± 34 |
| SAMT-A324M | n.d. | 5900 ± 260 | 980 ± 150 | 1700 ± 94 | n.d. | 290 ± 49 |
| SAMT-A324T | 260 ± 13 | 9600 ± 240 | n.d. | 420 ± 120 | n.d. | 1400 ± 6.4 |
| SAMT-F359H | n.d. | 690 ± 150 | n.d. | n.d. | n.d. | n.d. |

Mutant enzymes were kinetically characterized to compare their activity profiles and Km values with AA, SA, and BA to the wild-type CsSAMT (Table 1). While the wild-type CsSAMT had a low Km value of 105 μM for SA, all of the mutant enzymes had increased Km values relative to wild type. For the A324M and F359H mutants, activity was linear to 1 mM when SA concentrations were varied, meaning the Km value had increased to the point where it was unable to be saturated (Figure S8). These increased Km values suggest that all six residues are essential for SA recognition.

While all of the mutations increased Km values with SA, we were surprised that three of the mutants, C319S, M320I, and A324T had increased activity with 1 mM SA compared to the wild-type CsSAMT (Table 1). However, mutations at positions Met320 and Ala324 to different residues, Leu and Met, respectively, led to 35% less activity with SA compared to the wild-type enzyme, demonstrating that amino acid substitutions at these positions are key for SA activity. The F359H mutant had very low activity with SA, and the Km value was also not able to be captured, which indicates that having the aromatic, nonpolar Phe in the CsSAMT active site may be critical for binding aromatic substrates.

When the mutant enzymes were assayed with AA, four of the mutations, Q263C, Q263Y, C319S, and A324M, introduced detectable Km values when AA was varied (Table 1). The mutations at position 263 led to the lowest Km values of any of the mutants, 230 and 480 μM, when a tyrosine or cysteine was introduced, respectively. For both of these Gln263 mutants, the Km value for AA is reduced to almost half that of SA. The C319S mutant still had a lower Km value for SA (180 μM) than for AA (790 μM). For A324M, the Km value of 980 μM with AA was high, but AA was the only substrate for which the Km value was able to be captured for this mutant. Taken together, the lower Km values for AA relative to other substrates for Q263C, Q263Y, and A324M imply that these two residues (Gln263 and Ala324) are critical for AA recognition in CsSAMT.

Six of the mutations (Q263C, Q263Y, C319S, M320I, M320L, and A324M) increased activity with AA relative to the wild-type enzyme, ranging from 1.4-fold higher activity for the M320L mutant to 3.3-fold higher activity for the Q263Y mutant (Table 1). When we compared the ratio of activity of AA to SA for each of the mutants relative to that of the wild-type CsSAMT, these six mutants also all had increased ratios (Figure S9). The Q263Y mutant had a ratio of AA to SA that was 3.9-fold higher than that of the wild-type enzyme, which represents a substantial shift in preference. The AA to SA ratio for the Q263C and A324M mutants was 2.5-fold higher than that of the wild-type enzyme, highlighting the importance of these two positions for conferring AA activity.

When the mutants were assayed with BA, none of the mutations led to a detectable Km value (Table 1). Only one mutation, M320I, led to 38% increased activity with BA compared to the wild-type enzyme. Notably, the Q263Y, M320L, and A324M mutants, which had high activity with AA, had 12.7-, 6.1-, and 6.6-fold lower activity with BA, respectively, further emphasizing the role of these positions in AA substrate specificity.

Overall, we have identified three residues in the CsSAMT active site that confer AA activity: Gln263, Cys319, and Ala324. One or more mutations at these positions reduced the Km value with AA and also increased the ratio of activity with AA to SA relative to the wild-type enzyme (Table 1 and Figure S9). While the CsSAMT has high activity with AA, it is possible that additional SABATH methyltransferases that have yet to be characterized may also synthesize MeAA. There may also be an AMAT in *Citrus* spp. that is responsible for generating anthranilate esters using anthraniloyl-CoA and methanol, similar to the enzyme in grapes (Wang and De Luca, 2005, Yang *et al*., 2020).

### Revisiting substrate specificity of the maize AAMT1

Although the maize AAMT1 had already been probed for SA specificity, we wondered whether introducing residues that were found to be critical for substrate recognition in the *Citrus sinensis* SAMT would impart SA activity in ZmAAMT1. To test this, we compared the active site of the ZmAAMT1 enzyme to other SABATH methyltransferases, including the CsSAMT and the grape enzymes, 4g02123 and 4g02169, that were identified in this study (Figure 3). Twelve single mutations were introduced— F164S, F164Y, C165S, Q167W, Y246H, L329M, M333A, M333T, H368F, H368T, and H368Y, as well as the Y246W mutant that was previously shown to introduce activity with SA (Kollner *et al*., 2010). The resulting mutant proteins were inactive in two *in vitro* assays with the exception of the ZmAAMT1 L329M mutant, which had a 3.8-fold higher AA activity when assayed using ^14^C-labeled SAM and 25% higher activity when MeAA production was quantified by GC-MS (Figure 4a; Figure S10 and S11). Köllner and colleagues had success in assaying Y246W, Q167M, and Q167H mutations in the same enzyme and found that they were active with AA, and the Y246W mutant introduced 13% activity with SA relative to activity with AA (Kollner *et al*., 2010). While we did not see activity with these mutants *in vitro*, we did detect MeAA in the media of *E. coli* that were expressing each mutant gene when AA was added, suggesting that all of the mutant enzymes were indeed active (Figure S12).

**Figure 4.**
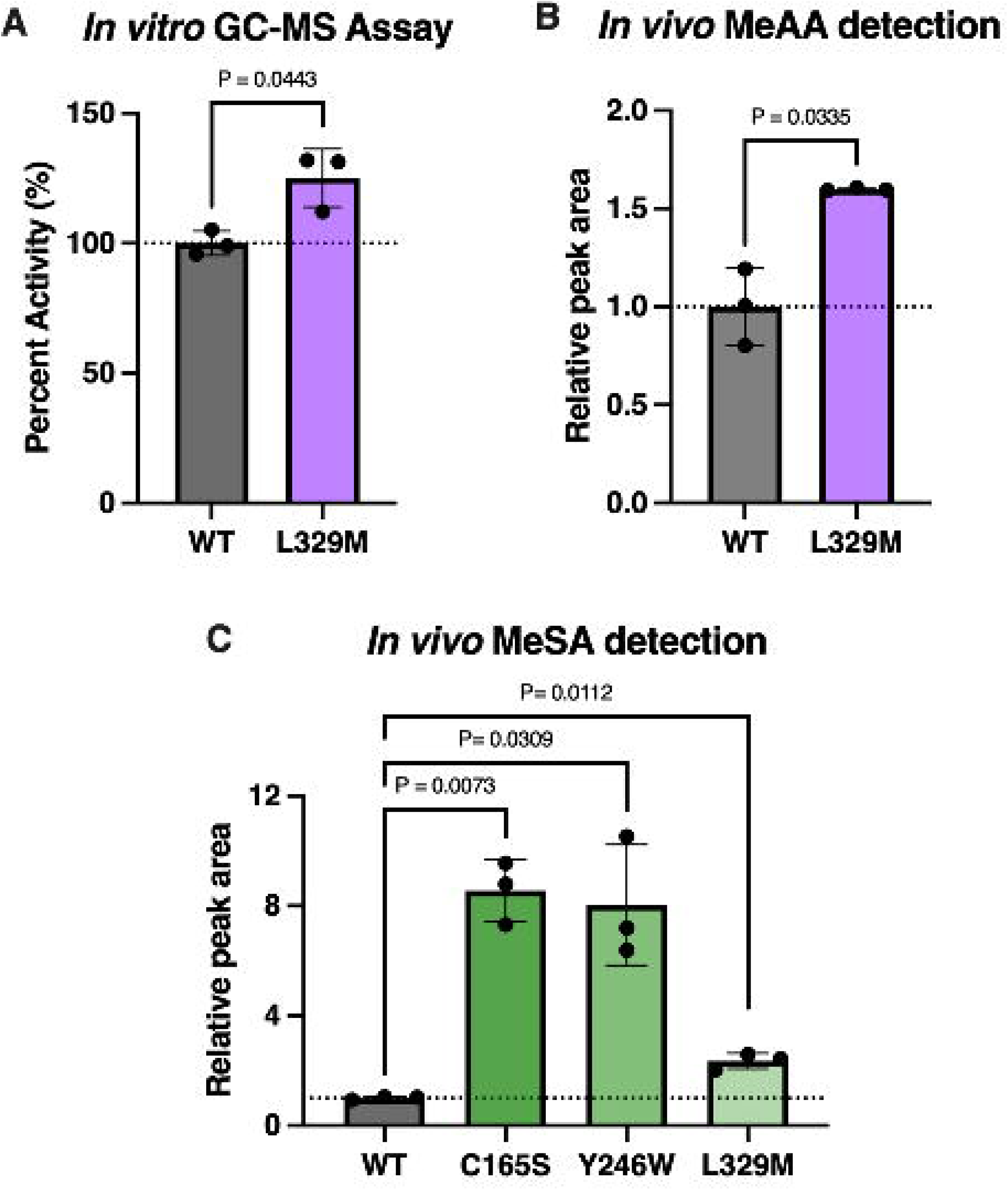
Identification of residues in the ZmAAMT1 active site that impart activity with MeSA. (A) The increased activity of the L329M mutant compared to the wild-type ZmAAMT1 was confirmed by quantitative GC-MS. (B) MeAA was compared in *E. coli* cultures expressing *ZmAAMT1* or the *L329M* mutant, which was determined by GC-MS. (C) MeSA was compared in *E. coli* cultures expressing *ZmAAMT1* or the *C165S, Y246W*, or *L329M* mutants, which was determined by GC-MS. For all points, n=3; where error bars are not visible, standard deviation is too small to visualize. P-values were calculated using a two-tailed Welch’s t-test. For A, there was no statistically significant difference (α level of 0.05).

Because our aim was to understand how the ZmAAMT1 has such high specificity for AA in comparison to SA, we also included one additional ZmAAMT1 mutant, I274Q, based on the data from the CsSAMT Q263Y mutant (Figure 3 and Table 1). The activity of the wild-type and each of the thirteen mutant enzymes was determined *in vivo* by adding SA to the *E. coli* cultures that were expressing each gene, and extracts of the spent media were analyzed by GC-MS for MeSA. Three ZmAAMT1 mutants— C165S, Y246W, and L329M—generated 8.5-, 8-, and 2-fold more MeSA, respectively, relative to the wild-type enzyme when expressed *in vivo* (Figure 4c). We did not see increased MeSA generated by the other ten mutants *in vivo* (Figure S13). Taken together, these results demonstrate that Cys165 and Leu329 also contribute to AA specificity in ZmAAMT1, since mutating these residues to a Ser and Met, respectively, increased activity with SA (Figure 4c). Because the L329M mutant has overall increased activity with AA *in vitro* and *in vivo*, this has implications for production of MeAA in engineered organisms (Figure 4a and 4b)(Luo *et al*., 2019).

### Structure-function-sequence evolution across AAMTs

To begin to understand the evolutionary relationship of the AA-using enzymes investigated here—the grape methyltransferases (4g02122, 4g02123, 4g02169), the citrus SAMT, and the maize AAMT1, a phylogeny was constructed (Figure S14). The CsSAMT and ZmAAMT1 have already been phylogenetically characterized, and the dendrogram generated here is consistent with previous analyses (Köllner *et al*., 2010, Huang *et al*., 2016, Pollier *et al*., 2019, Dubs *et al*., 2022). Of the grape enzymes, two (4g02122 and 4g02123) share an ancestor with each other and with other SAMTs, and because these two genes are clustered in close proximity on chromosome 4 and are active with either AA or SA, the AAMT and N-MeAAMT activity in these two *Vitis* genes may have evolved through gene duplication followed by neofunctionalization. Interestingly, the third grape enzyme, 4g02169, clustered with characterized jasmonic acid methyltransferases (JMTs). This was also true of the strawberry (*Fragaria* spp.) AAMT, which also has shared ancestry with JMTs (Pillet *et al*., 2017). When we assayed Vv4g02169 with JA, there was activity, although it was 5-fold lower than the enzyme’s activity with AA (Figure 2a and S15).

To connect protein structure and function to sequence evolution, we compared the active site residues that were investigated here across all of the methyltransferases in the phylogeny (Figure S14). In the CsSAMT active site, we identified four residues that are important for conferring AA activity: Gln263, Cys319, and Ala324 (Figure 5a and Table 1). Gln263 is conserved in Vv4g02122, Vv4g02123, and the soybean AAMT, but variable across the other functionally characterized AAMTs (Tyr in Vv4g02169; Ile in ZmAAMT1; Asn in the alfalfa AAMT) (Figure 5c and S14). A notable trend is that enzymes that have a Gln in this position can all also methylate SA, but the AA-using enzymes that have a different residue have very little to no activity with SA (Table 1 and Figure 2a) (Pollier *et al*., 2019). The CsSAMT Cys319 residue is a Leu in Vv4g02122 and 4g02123 and a Ser in Vv4g02169; introducing a polar Ser at this position in the CsSAMT increased overall activity and increased activity with AA, suggesting that this position may contribute to the high AA activity in Vv4g02169 (Figure 5c and Table 1). The CsSAMT A324M mutation increased AA activity 2-fold, and this mutant only had a detectable Km value for AA. This Ala is conserved in most all AA-using methyltransferases but is a conserved Met in the maize AAMTs and a Gln in Vv4g02169 (Figure 5c and S14). Introducing a larger, nonpolar Met at this active site position may facilitate the positioning of the aromatic ring of AA, SA, and BA.

**Figure 5.**
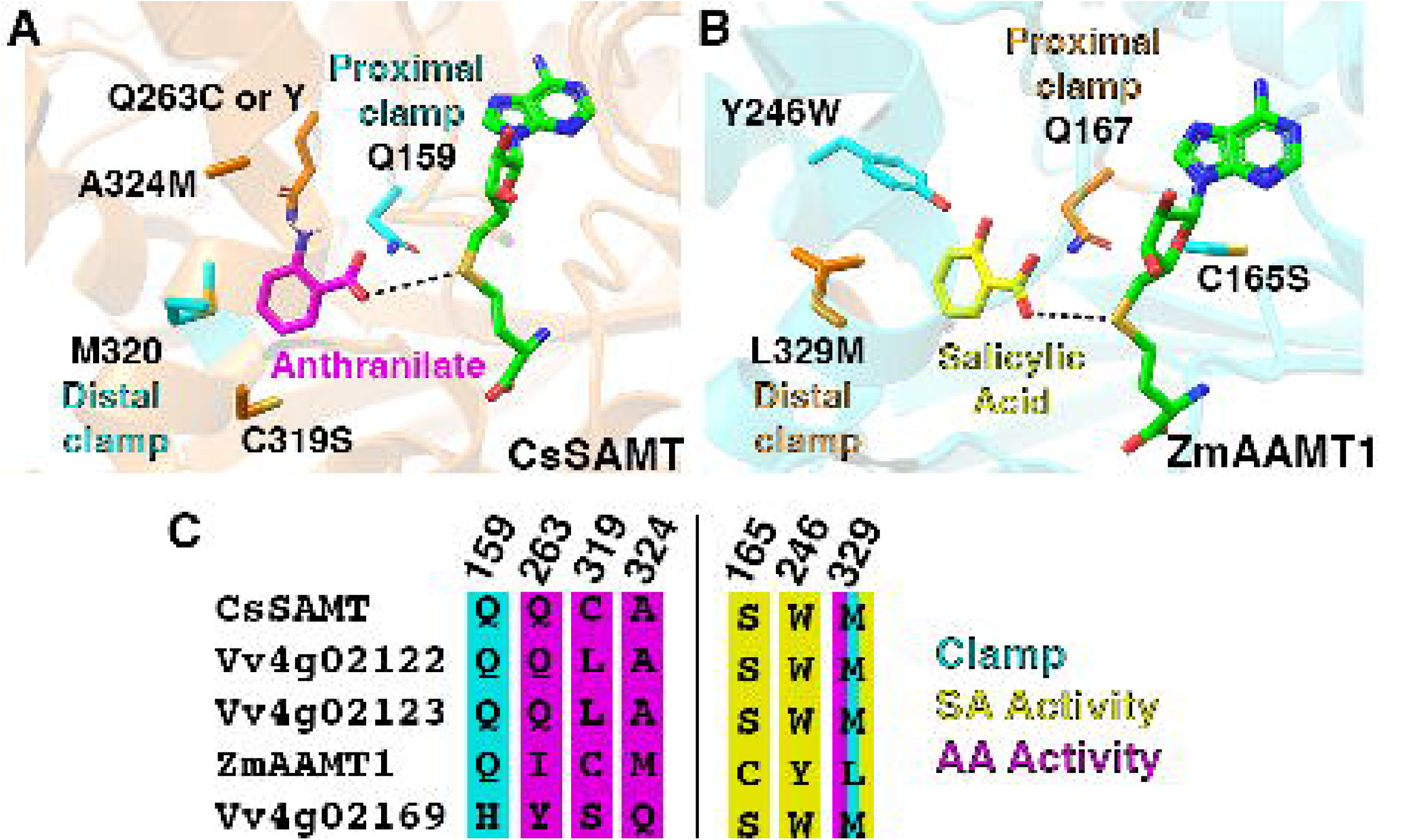
Connecting substrate specificity to protein structure. (A) The *Citrus sinensis* SAMT active site with anthranilate (magenta) and SAH (green) bound. Four mutations at three residues (Q263C, Q263Y, C319S, and A324M) were important for conferring AA activity. Gln159 and Met320 (cyan) form a molecular clamp. (B) The *Zea mays* AAMT1 active site with salicylic acid (yellow) and SAH (green) bound. Three mutations (C165S, Y246W, and L329M) increased activity with SA; L329M also increased activity with AA. Gln167 and Leu329 (orange) form a molecular clamp. (C) Amino acid sequence alignment of the active site residues that conferred AA activity in CsSAMT (left of the black bar; numbers correspond to CsSAMT) or that conferred SA activity in ZmAAMT1 (right; numbers correspond to ZmAAMT1).

In the maize AAMT1 active site, we identified two residues—in addition to Tyr246 that had been identified previously (Kollner *et al*., 2010)—that are important for conferring SA activity: Cys165 and Leu329 (Figure 4c and 5b). Cys165 is a polar Ser in most all of the other AA and SA methyltransferases (Figure 5c and S14). Having a polar residue at this position may aid in positioning acyl acid substrates for the methyl group transfer from SAM. The maize AAMT Leu329 (which corresponds to position 320 in the CsSAMT) is a nonpolar Met or Ile in the other AAMTs, so the fact that the conservative L329M mutation led to increased MeAA and MeSA production was surprising. Because Trp levels in maize are low (Kaur *et al*., 2020), having a Leu in position 329 that reduces AAMT1 activity suggests that there is weaker selective pressure for MeAA biosynthesis in maize (Bar-Even *et al*., 2011). Also, the reduced ZmAAMT1 activity may ensure that there is sufficient AA to support the biosynthesis of Trp, benzoxazinoids, and auxin hormones (Maeda and Dudareva, 2012).

Other investigations of active site residues in SAMTs have identified two Met residues that form a molecular clamp in the *Clarkia breweri* SAMT to position the benzyl ring of SA in the active site (Zubieta *et al*., 2003). This clamp is formed by Gln159 and Met320 in CsSAMT; Gln167 and Leu329 in ZmAAMT1; Gln156 and Met314 in Vv4g02123; and His156 and Met316 in Vv4g02169 (Figure 5). The proximal clamp residue has also been implicated in substrate preference for SA versus BA, and others have found that having a Met at position 150 is a key determinant of SA substrate preference (Barkman *et al*., 2007, Han *et al*., 2018, Dubs *et al*., 2022). In our experiments, the proximal clamp residue seemed to matter less for determining AA substrate specificity, which is expected given that this site is ancestral in SAMT enzymes (Dubs *et al*., 2022). When the CsSAMT Gln159 was mutated to a His, activity was reduced with all three acyl acid substrates—SA, AA, and BA (Table 1). The distal clamp residue seemed to impact substrate specificity more. The ZmAAMT1 L329M mutant in effect introduces the same clamp residues (Gln and Met) that are found in the CsSAMT and Vv4g02123 active sites, both of which have high activity with SA (Table 1 and Figure 2d and 2g). The fact that ZmAAMT1 L329M has higher activity overall further suggests that having the thioether moiety in the distal clamp position is important for AA and SA activity (Figure 4).

Overall, our findings, coupled with those from other SAM-dependent carboxy methyltransferases, highlights that there are multiple active site residues that confer substrate specificity. There is likely some context-dependence as well, meaning that the combination of residues that make up the active site collectively impart specificity for AA versus other acyl acids. Additionally, we found residues outside the active site in the ‘Concord’ 4g02123 that altered activity, which was unanticipated (Figure 2e-g and 2i). As new AAMTs are discovered and more sequence data are available, it may become easier to predict which combinations of residues are required for AA activity.

### Conclusions and Future Directions

Here we identified two *Vitis vinifera* SAM-dependent methyltransferases (Vv4g02123 and Vv4g02169), as well as a ‘Concord’ grape ortholog of 4g02123, that can generate the volatile MeAA (Table 1 and Figure 2). Probing the molecular basis of AA recognition in a citrus SAMT led us to identify three key residues responsible for AA activity: Gln263, Cys319, and Ala324 (Table 1 and Figure 5). Conversely, in the maize AAMT1, examining the basis of SA activity led us to identify two residues, Cys165 and Leu329, in addition to the previously identified Tyr246 (Kollner *et al*., 2010). MeAA has been under sampled in plants, especially in below-ground tissues (Pollier *et al*., 2019). Since all plants synthesize AA as an intermediate in Trp biosynthesis (Li *et al*., 2023), there may be many more plant methyltransferases that can act on AA that remain to be discovered.

Because MeAA is commercially relevant, there is interest in using engineered microorganisms for MeAA production, as opposed to current practices that use non-renewable petroleum sources (Yadav and Krishnan, 1998). MeAA synthesized from yeast or bacteria is considered a natural flavor, and an engineered bacterium strain overexpressing the maize *AAMT1* gene generated 5.74 g/L of MeAA (Luo *et al*., 2019). The *Vitis* enzymes identified in our experiments may promote even higher yield in a synthetic biology platform. Identification of a DiMeAA-producing enzyme may also have implications for human health, as evidence suggests that DiMeAA has anti-nociceptive, anti-depression, and anti-anxiety effects in mice (Radulovic *et al*., 2013, Pinheiro *et al*., 2014). This is unsurprising considering a number of anthranilate derived alkaloids are bioactive (Shende *et al*., 2024). Additionally, identifying grape AAMTs has the potential to improve grape breeding to decrease MeAA levels in wine and eliminate the ‘foxy’ odor. Our findings highlight the power of coupling existing ‘omics’ data with characterizations of enzyme structure and function to reveal potential avenues to engineer plant volatile biosynthesis.

## EXPERIMENTAL PROCEDURES

### Protein visualization and molecular docking

Structural models of all proteins were obtained from AlphaFold (Jumper *et al*., 2021) (Uniprot IDs: CsSAMT: A0A1S8AD96; Vv4g02123: A0A438BRB9; Vv4g02169: D7SNV7; and ZmAAMT1: D9J0Z7), and anthranilate was docked into each active site using AutoDock Vina (ver. 1.1.2) with a grid box of 40 x 40 x 40 Å and the exhaustiveness set to 8 (Trott and Olson, 2010, Forli *et al*., 2016). The results were visualized in PyMOL (ver. 2.5.7) (https://www.pymol.org/2/). Dimer complexes were generated using ColabFold v1.5.5 (Mirdita *et al*., 2022), and SAH was added to the dimers using AlphaFill (Hekkelman *et al*., 2023).

### Site-directed mutagenesis

The wild-type amino acid sequences for CsSAMT (XP_006466836.1), ZmAAMT1 (D9J0Z8.1), Vv4g02122 (XP_002262759.2), Vv4g02123 (XP_002262676.1), Vv4g02169 (XP_002263459.1), Vv12g00725 (XP_002267308.1) were used for codon-optimized gene synthesis (Data S1) and cloning into the *E. coli* pET28a expression vector by Twist Biosciences. The wild-type plasmids were used as a template for site-directed mutagenesis PCR, and mutations were introduced using the QuikChange PCR method (Agilent). Oligonucleotides were designed using PrimerX (https://www.bioinformatics.org/primerx/) or the Agilent QuikChange Primer Design tool (Table S1). The successful introduction of each mutation was determined by Sanger sequencing (Genewiz).

### Protein expression and purification

Each construct was transformed into *E. coli* BL21 (DE3) cells (New England Biolabs). Cells were cultured in one liter of Terrific broth until the *A*_600nm_ reached 0.6–0.8, at which time the incubator temperature was lowered to 16°C and protein expression was induced using 1 mM IPTG. After overnight incubation, cells were pelleted by centrifugation (5000*g*; 15 min) and resuspended in 40 mL of lysis buffer (50 mM Tris (pH 8.0), 500 mM NaCl, 20 mM imidazole, 10% glycerol, and 1% Tween-20). Following sonication, cell debris was removed by centrifugation (13,000*g*; 60 min) and the resulting lysate was passed over a Ni^2+^- nitrilotriacetic acid column (1.5 x 12 cm) equilibrated in the lysis buffer. The column was then washed (50 mM Tris (pH 8.0), 500 mM NaCl, 20 mM imidazole, and 10% glycerol) and bound His-tagged protein eluted (50 mM Tris (pH 8.0), 500 mM NaCl, 250 mM imidazole, and 10% glycerol). Protein aliquots were stored at -80°C. Protein concentration was determined by the Bradford method (Bio-Rad) with bovine serum albumin as the standard.

### Luminescence Kinetics Assay

All *Citrus* and *Vitis* enzymes were assayed using the MTase-Glo™ methyltransferase assay (Promega) (Hsiao *et al*., 2016). A 4x reaction buffer was prepared so that the final concentrations in each well were 20 mM Tris (pH 8.0), 50 mM NaCl, 1 mM EDTA, 3 mM MgCl_2_, 200 μg/mL bovine serum albumin, and 1 mM TCEP. The concentration of each protein was adjusted to 14-22 μg, and the wild-type and mutant enzymes were each assayed in triplicate with SAM (100 μM for CsSAMT and its mutants; 20 μM for 4g02123 from *V. vinifera* and its ortholog from ‘Concord’; 150 μM for Vv4g02122 and Vv4g02169) and 0-1 mM substrate with at least 5 concentrations of AA, SA, BA, or N-MeAA (Figure S16). The 20 μL reactions were initiated by the addition of substrates, and the reactions were incubated at 37°C for 30 minutes. Reactions were quenched by the addition of 5 μL of 0.5% trifluoroacetic acid before adding the 5 μL of 6x MTase-Glo reagent and subsequent 30 μL of MTase-Glo detection solution. Luminescence readings were measured using a BioTek Cytation 1 plate reader. The enzyme activity was calculated in terms of nmol of product per milligram of enzyme using a SAH standard curve ranging from 0 to 12,500 μM with five concentrations in between (n=3) (Figure S1).

### *In vitro* assays for MeAA detection

A 100 μL assay (100 mM HEPES (pH 7.5), 2 mM EDTA (pH 8.0), 10% glycerol, 25 μg of methyltransferase, and 1 mM anthranilate or salicylic acid) was initiated by the addition of a final concentration of 2 mM SAM, and 1 mL of hexanes was overlaid on top of the reaction to trap the volatile products. The reaction mixture was then maintained at 37°C for 1 hour, after which time the reactions were vortexed to quench the reaction. For quantitative analysis, 550 nM ethyl anthranilate was added to the hexane as an internal standard for GC-MS analysis.

### *In vivo* assay for MeAA and MeSA detection

Following the methods of Dubs and colleagues (Dubs *et al*., 2022), cultures of *E. coli* BL21 harboring the pET28a plasmid with the *ZmAAMT1* gene, the ‘Concord’ ortholog of *4g02169*, or their mutants were grown in 50 mL of Terrific broth until the *A*_600nm_ reached 0.6–0.8, at which time the incubator temperature was lowered to 16°C and protein expression was induced using 1 mM IPTG. At the same time, salicylic acid or anthranilate were added to the cultures at a final concentration of 200 μM. The following day, cells were pelleted (3500*g* for 10 minutes), and 5 mL of hexanes were added to 45 mL of the spent media. After vortex mixing and centrifuging (3500*g* for 5 minutes), the hexane layer was analyzed by GC-MS.

### GC-MS Analysis

The ZmAAMT1 samples were analyzed using a 7820A gas chromatograph system coupled to a 5977B mass selective detector (Agilent Technologies). Data were acquired and processed using MassHunter Agilent Technologies™ Software (Santa Clara, CA, USA). The selected ion monitoring mode was used to search for specific ions to identify and quantify O-methyl anthranilate (m/z of 92, 119 and 151) and O-ethyl anthranilate and dimethyl anthranilate (m/z of 92, 119, 151, and 165), and external standards of pure compounds were included with each run. Each ion was measured with a dwell time of 100 ms. A solvent delay for the MS of 2.0 min was used. A splitless injection volume of 1 μL was injected onto an HP-5 column with 5% phenyl methylpolysiloxane stationary phase (Agilent) with dimensions of 30 m x 0.25 mm x 0.25 μm. Helium was used as the carrier gas with a flow rate of 1.2 mL/min. The injection port was set at 280°C with a purge flow of 50 mL/min at 0.4 min. For the ZmAAMT1 *in vitro* assay samples, oven temperature was initially set at 110°C, held at that temperature for 2 minutes and then increased at a rate of 30°C/min until 270°C, where it was held for 2 minutes; total run time was 9.33 min. For the *in vivo E. coli* extracts and the *in vitro* grape assays, the oven temperature was initially set at 60°C, held for 2 minutes and then increased at a rate of 30°C/min until 300°C, where it was held for 2 minutes; total run time was 12 min. Data were acquired and processed using MassHunter Agilent Technologies™ Software. For MeAA quantification produced by ZmAAMT1 *in vitro*, the limit of detection and limit of quantification were determined, and a calibration curve was generated using triplicate values from 0-2000 nM for MeAA and the internal standard EtAA (Figure S11) (Hubaux and Vos, 1970). The products of the *Vitis* enzymes were confirmed *in vitro* using the same method on a 7890B GC coupled to a 7000C triple-quadrupole mass spectrometer (Agilent Technologies).

### Assay for ZmAAMT1 activity using^14^C-SAM

The assay followed a published method (Kollner *et al*., 2010), and each 100 μL reaction contained 25 μg of methyltransferase buffered with 100 mM HEPES (pH 7.5), 2 mM EDTA (pH 8.0), and 10% glycerol. Anthranilate and salicylic acid were tested at a final concentration of 1 mM, and the reactions were initiated by the addition of 3 μL of 52.6 mCi/mmol S-[methyl-^14^C]adenosyl-L-methionine (PerkinElmer). The reaction was overlaid with 1 mL of pentane and incubated 37°C for 1 hour. The reactions were quenched by vortexing, and the pentane layer was mixed 1:5 (v/v) with Scintiverse BD Biodegradable LSC cocktail. The scintillation vials were read in a Beckman LS 6500 multiple-purpose scintillation counter (96.38% efficiency). Counts per minute were used to calculate the activity in nmol/mg of protein.

### Phylogenetic Analysis

Homologous sequences and an outgroup were selected from SAMT and JMT clades based on comprehensive characterization and phylogenetic analyses of plant SABATH methyltransferases (Kollner *et al*., 2010, Koeduka *et al*., 2020, Dubs *et al*., 2022). Amino acid sequences were aligned using MAFFT version 7 (Katoh and Standley, 2013). Protein phylogenies were inferred using the IQ-TREE web server with default parameters and 10,000 ultrafast bootstrap replicates (Nguyen *et al*., 2014, Hoang *et al*., 2017, Kalyaanamoorthy *et al*., 2017). The best fit model according to Bayesian Information Criterion was the JTT model with a discrete Gamma model with 4 rate categories (Jones *et al*., 1992, Yang, 1994). The dendrogram was visualized using FigTree (version 1.4.4).

## AUTHOR CONTRIBUTIONS

N.C., M.D, M.A.F., C.K.H., H.T., and A.P.W. designed the research; M.D, M.A.F., C.F., C.K.H., H.T., and A.P.W. performed the research; all authors contributed to data analysis; C.K.H. wrote the paper with all authors providing editorial input.

## ACKNOWLEDGEMENTS

We thank Holland lab members for their feedback on this work, as well as Drs. Lucas Busta, Tara Enders, and Thomas Smith for helpful conversations. This research was supported by funds from Williams College and the U.S. National Science Foundation (MCB-2214883 to C.K.H.), and H.T. was funded by the American Society of Plant Biologists through a Summer Undergraduate Research Fellowship. Portions of the mass spectral data were obtained at the University of Massachusetts Mass Spectrometry Core Facility, RRID:SCR_019063.

## SUPPORTING INFORMATION

### Supporting figures

**Figure S1.** Kinetic plots of the activities of *Vitis vinifera* (Vv) 4g02122, 4g02123, and ‘Concord’ ortholog of 4g02123 with BA or SA.

**Figure S2.** GC-MS data confirms that 4g02123 from ‘Concord’ and wine grapes (Vv) synthesizes methyl anthranilate (MeAA) *in vitro*.

**Figure S3.** GC-MS data confirms that 4g02123 from ‘Concord’ and wine grapes (Vv) synthesizes dimethyl anthranilate (DiMeAA) *in vitro*.

**Figure S4.** Promoter sequence comparisons for *Vitis vinifera* Vv4g02123 and its orthologs in *Vitis labrusca* and ‘Concord’ grapes.

**Figure S5.** Promoter sequence comparisons for *Vitis vinifera* Vv4g02169 and its orthologs in *Vitis labrusca* and ‘Concord’ grapes.

**Figure S6.** Coding sequence alignment of *Vitis vinifera* Vv4g021**2**3 and its orthologs in *Vitis labrusca* and ‘Concord’ grapes.

**Figure S7.** Coding sequence alignment of *Vitis vinifera* Vv4g02169 and its orthologs in *Vitis labrusca* and ‘Concord’ grapes.

**Figure S8.** Michaelis-Menten plots of *Citrus sinensis* (Cs) SAMT activity with SA, AA or BA.

**Figure S9.** Ratio of activity with AA relative to SA for the *Citrus sinensis* SAMT.

**Figure S10.** Percent activity of ZmAAMT1 mutants with 1 mM anthranilate or 1 mM salicylic acid relative to wild type, which was determined using ^14^C-SAM.

**Figure S11.** GC-MS data for *in vitro* MeAA quantification for the wild-type ZmAAMT1 enzyme and the L329M mutant.

**Figure S12.** GC-MS chromatograms for *in vivo* ZmAAMT1 MeAA detection.

**Figure S13.** GC-MS chromatograms for *in vivo* ZmAAMT1 MeSA detection.

**Figure S14.** Dendrogram of plant acyl acid methyltransferases.

**Figure S15.** Jasmonic acid (JA) activity in the Vv4g02169 enzyme.

**Figure S16.** SAM kinetics plots of CsSAMT with saturating salicylic acid and Vv4g02122, Vv4g02123 and Vv4g02169 with saturating anthranilate.

### Supporting tables

**Table S1**. List of primers used for site-directed mutagenesis.

### Supporting data

**Data S1.** Codon-optimized gene sequences of *AAMTs* and *SAMTs*.

## REFERENCES

Avery, M.L., Decker, D.G., Humphrey, J.S., Aronov, E., Linscombe, S.D. and Way, M. (1995) Methyl anthranilate as a rice seed treatment to deter birds. The Journal of wildlife management, 50-56.

Azam, M., Song, M., Fan, F., Zhang, B., Xu, Y., Xu, C. and Chen, K. (2013) Comparative analysis of flower volatiles from nine citrus at three blooming stages. Int J Mol Sci, 14, 22346–22367.

Baldwin, I.T. (2010) Plant volatiles. Curr Biol, 20, R392-397.

Bar-Even, A., Noor, E., Savir, Y., Liebermeister, W., Davidi, D., Tawfik, D.S. and Milo, R. (2011) The moderately efficient enzyme: evolutionary and physicochemical trends shaping enzyme parameters. Biochemistry, 50, 4402–4410.

Barkman, T.J., Martins, T.R., Sutton, E. and Stout, J.T. (2007) Positive Selection for Single Amino Acid Change Promotes Substrate Discrimination of a Plant Volatile-Producing Enzyme. Molecular Biology and Evolution, 24, 1320–1329.

Bracker, L.B., Gong, X., Schmid, C., Dawid, C., Ulrich, D., Phung, T., Leonhard, A., Ainsworth, J., Olbricht, K., Parniske, M. and Gompel, N. (2020) A strawberry accession with elevated methyl anthranilate fruit concentration is naturally resistant to the pest fly Drosophila suzukii. PLoS One, 15, e0234040.

Dubs, N.M., Davis, B.R., de Brito, V., Colebrook, K.C., Tiefel, I.J., Nakayama, M.B., Huang, R., Ledvina, A.E., Hack, S.J., Inkelaar, B., Martins, T.R., Aartila, S.M., Albritton, K.S., Almuhanna, S., Arnoldi, R.J., Austin, C.K., Battle, A.C., Begeman, G.R., Bickings, C.M., Bradfield, J.T., Branch, E.C., Conti, E.P., Cooley, B., Dotson, N.M., Evans, C.J., Fries, A.S., Gilbert, I.G., Hillier, W.D., Huang, P., Hyde, K.W., Jevtovic, F., Johnson, M.C., Keeler, J.L., Lam, A., Leach, K.M., Livsey, J.D., Lo, J.T., Loney, K.R., Martin, N.W., Mazahem, A.S., Mokris, A.N., Nichols, D.M., Ojha, R., Okorafor, N.N., Paris, J.R., Reboucas, T.F., Sant’Anna, P.B., Seitz, M.R., Seymour, N.R., Slaski, L.K., Stemaly, S.O., Ulrich, B.R., Van Meter, E.N., Young, M.L. and Barkman, T.J. (2022) A Collaborative Classroom Investigation of the Evolution of SABATH Methyltransferase Substrate Preference Shifts over 120 My of Flowering Plant History. Mol Biol Evol, 39.

Dudareva, N., Negre, F., Nagegowda, D.A. and Orlova, I. (2006) Plant Volatiles: Recent Advances and Future Perspectives. Critical Reviews in Plant Sciences, 25, 417–440.

Dudareva, N., Pichersky, E. and Gershenzon, J. (2004) Biochemistry of plant volatiles. Plant physiology, 135, 1893–1902.

Effmert, U., Saschenbrecker, S., Ross, J., Negre, F., Fraser, C.M., Noel, J.P., Dudareva, N. and Piechulla, B. (2005) Floral benzenoid carboxyl methyltransferases: from in vitro to in planta function. Phytochemistry, 66, 1211–1230.

Forli, S., Huey, R., Pique, M.E., Sanner, M.F., Goodsell, D.S. and Olson, A.J. (2016) Computational protein-ligand docking and virtual drug screening with the AutoDock suite. Nat Protoc, 11, 905–919.

Gong, Q., Wang, Y., He, L., Huang, F., Zhang, D., Wang, Y., Wei, X., Han, M., Deng, H., Luo, L., Cui, F., Hong, Y. and Liu, Y. (2023) Molecular basis of methyl-salicylate-mediated plant airborne defence. Nature, 622, 139–148.

Gonzalez-Mas, M.C., Rambla, J.L., Lopez-Gresa, M.P., Blazquez, M.A. and Granell, A. (2019) Volatile Compounds in Citrus Essential Oils: A Comprehensive Review. Front Plant Sci, 10, 12.

Han, X.M., Yang, Q., Liu, Y.J., Yang, Z.L., Wang, X.R., Zeng, Q.Y. and Yang, H.L. (2018) Evolution and Function of the Populus SABATH Family Reveal That a Single Amino Acid Change Results in a Substrate Switch. Plant Cell Physiol, 59, 392–403.

Hekkelman, M.L., de Vries, I., Joosten, R.P. and Perrakis, A. (2023) AlphaFill: enriching AlphaFold models with ligands and cofactors. Nature Methods, 20, 205–213.

Hoang, D.T., Chernomor, O., von Haeseler, A., Minh, B.Q. and Vinh, L.S. (2017) UFBoot2: Improving the Ultrafast Bootstrap Approximation. Molecular Biology and Evolution, 35, 518–522.

Hsiao, K., Zegzouti, H. and Goueli, S.A. (2016) Methyltransferase-Glo: a universal, bioluminescent and homogenous assay for monitoring all classes of methyltransferases. Epigenomics, 8, 321–339.

Huang, R., O’Donnell, A.J., Barboline, J.J. and Barkman, T.J. (2016) Convergent evolution of caffeine in plants by co-option of exapted ancestral enzymes. Proc Natl Acad Sci U S A, 113, 10613–10618.

Hubaux, A. and Vos, G. (1970) Decision and detection limits for calibration curves. Analytical Chemistry, 42, 849–855.

Jones, D.T., Taylor, W.R. and Thornton, J.M. (1992) The rapid generation of mutation data matrices from protein sequences. Bioinformatics, 8, 275–282.

Jumper, J., Evans, R., Pritzel, A., Green, T., Figurnov, M., Ronneberger, O., Tunyasuvunakool, K., Bates, R., Žídek, A., Potapenko, A., Bridgland, A., Meyer, C., Kohl, S.A.A., Ballard, A.J., Cowie, A., Romera-Paredes, B., Nikolov, S., Jain, R., Adler, J., Back, T., Petersen, S., Reiman, D., Clancy, E., Zielinski, M., Steinegger, M., Pacholska, M., Berghammer, T., Bodenstein, S., Silver, D., Vinyals, O., Senior, A.W., Kavukcuoglu, K., Kohli, P. and Hassabis, D. (2021) Highly accurate protein structure prediction with AlphaFold. Nature, 596, 583–589.

Kalyaanamoorthy, S., Minh, B.Q., Wong, T.K.F., von Haeseler, A. and Jermiin, L.S. (2017) ModelFinder: fast model selection for accurate phylogenetic estimates. Nature Methods, 14, 587–589.

Katoh, K. and Standley, D.M. (2013) MAFFT multiple sequence alignment software version 7: improvements in performance and usability. Mol Biol Evol, 30, 772–780.

Kaur, R., Kaur, G., Vikal, Y., Gill, G.K., Sharma, S., Singh, J., Dhariwal, G.K., Gulati, A., Kaur, A. and Kumar, A. (2020) Genetic enhancement of essential amino acids for nutritional enrichment of maize protein quality through marker assisted selection. Physiology and Molecular Biology of Plants, 26, 2243–2254.

Knudsen, J.T., Eriksson, R., Gershenzon, J. and Ståhl, B. (2006) Diversity and distribution of floral scent. The botanical review, 72, 1–120.

Koeduka, T., Suzuki, H., Taguchi, G. and Matsui, K. (2020) Biochemical characterization of the jasmonic acid methyltransferase gene from wasabi (Eutrema japonicum). Plant Biotechnol (Tokyo), 37, 389–392.

Kollner, T.G., Lenk, C., Zhao, N., Seidl-Adams, I., Gershenzon, J., Chen, F. and Degenhardt, J. (2010) Herbivore-induced SABATH methyltransferases of maize that methylate anthranilic acid using s-adenosyl-L-methionine. Plant Physiol, 153, 1795–1807.

Köllner, T.G., Lenk, C., Zhao, N., Seidl-Adams, I., Gershenzon, J., Chen, F. and Degenhardt, J. (2010) Herbivore-induced SABATH methyltransferases of maize that methylate anthranilic acid using s-adenosyl-L-methionine. Plant Physiol, 153, 1795–1807.

Li, M., Tadfie, H., Darnell, C.G. and Holland, C.K. (2023) Biochemical investigation of the tryptophan biosynthetic enzyme anthranilate phosphoribosyltransferase in plants. Journal of Biological Chemistry, 299.

Lin, J., Massonnet, M. and Cantu, D. (2019) The genetic basis of grape and wine aroma. Hortic Res, 6, 81.

Lin, J., Mazarei, M., Zhao, N., Zhu, J.J., Zhuang, X., Liu, W., Pantalone, V.R., Arelli, P.R., Stewart, C.N., Jr. and Chen, F. (2013) Overexpression of a soybean salicylic acid methyltransferase gene confers resistance to soybean cyst nematode. Plant Biotechnol J, 11, 1135–1145.

Luo, Z.W., Cho, J.S. and Lee, S.Y. (2019) Microbial production of methyl anthranilate, a grape flavor compound. Proc Natl Acad Sci U S A, 116, 10749–10756.

Maeda, H. and Dudareva, N. (2012) The shikimate pathway and aromatic amino acid biosynthesis in plants. Annual review of plant biology, 63, 73–105.

Mann, R.S., Ali, J.G., Hermann, S.L., Tiwari, S., Pelz-Stelinski, K.S., Alborn, H.T. and Stelinski, L.L. (2012) Induced release of a plant-defense volatile ‘deceptively’ attracts insect vectors to plants infected with a bacterial pathogen. PLoS Pathog, 8, e1002610.

Mason, J.R., Adams, M.A. and Clark, L. (1989) Anthranilate repellency to starlings: chemical correlates and sensory perception. The Journal of wildlife management, 55-64.

Mikiciuk, G., Chełpiński, P., Mikiciuk, M., Możdżer, E. and Telesiński, A. (2021) The Effect of Methyl Anthranilate-Based Repellent on Chemical Composition and Selected Physiological Parameters of Sweet Cherry (Prunus avium L.). Agronomy, 11, 256.

Mirdita, M., Schütze, K., Moriwaki, Y., Heo, L., Ovchinnikov, S. and Steinegger, M. (2022) ColabFold: making protein folding accessible to all. Nature Methods, 19, 679–682.

Nguyen, L.-T., Schmidt, H.A., von Haeseler, A. and Minh, B.Q. (2014) IQ-TREE: A Fast and Effective Stochastic Algorithm for Estimating Maximum-Likelihood Phylogenies. Molecular Biology and Evolution, 32, 268–274.

Pillet, J., Chambers, A.H., Barbey, C., Bao, Z., Plotto, A., Bai, J., Schwieterman, M., Johnson, T., Harrison, B., Whitaker, V.M., Colquhoun, T.A. and Folta, K.M. (2017) Identification of a methyltransferase catalyzing the final step of methyl anthranilate synthesis in cultivated strawberry. BMC Plant Biol, 17, 147.

Pinheiro, M.M.G., Radulović, N.S., Miltojević, A.B., Boylan, F. and Fernandes, P.D. (2014) Antinociceptive esters of N-methylanthranilic acid: Mechanism of action in heat-mediated pain. European Journal of Pharmacology, 727, 106–114.

Pollier, J., De Geyter, N., Moses, T., Boachon, B., Franco-Zorrilla, J.M., Bai, Y., Lacchini, E., Gholami, A., Vanden Bossche, R., Werck-Reichhart, D., Goormachtig, S. and Goossens, A. (2019) The MYB transcription factor Emission of Methyl Anthranilate 1 stimulates emission of methyl anthranilate from Medicago truncatula hairy roots. Plant J, 99, 637–654.

Radulovic, N.S., Miltojevic, A.B., Randjelovic, P.J., Stojanovic, N.M. and Boylan, F. (2013) Effects of methyl and isopropyl N-methylanthranilates from Choisya ternata Kunth (Rutaceae) on experimental anxiety and depression in mice. Phytother Res, 27, 1334–1338.

Rohde, B., Hans, J., Martens, S., Baumert, A., Hunziker, P. and Matern, U. (2008) Anthranilate N-methyltransferase, a branch-point enzyme of acridone biosynthesis. Plant J, 53, 541–553.

Sawler, J., Reisch, B., Aradhya, M.K., Prins, B., Zhong, G.-Y., Schwaninger, H., Simon, C., Buckler, E. and Myles, S. (2013) Genomics Assisted Ancestry Deconvolution in Grape. PLOS ONE, 8, e80791.

Schuman, M.C. (2023) Where, When, and Why Do Plant Volatiles Mediate Ecological Signaling? The Answer Is Blowing in the Wind. Annu Rev Plant Biol, 74, 609–633.

Shende, V.V., Bauman, K.D. and Moore, B.S. (2024) The shikimate pathway: gateway to metabolic diversity. Nat Prod Rep, 41, 604–648.

Simpson, M., Gurr, G.M., Simmons, A.T., Wratten, S.D., James, D.G., Leeson, G. and Nicol, H.I. (2011) Insect attraction to synthetic herbivore-induced plant volatile-treated field crops. Agricultural and Forest Entomology, 13, 45–57.

Sun, Q., Gates, M.J., Lavin, E.H., Acree, T.E. and Sacks, G.L. (2011) Comparison of Odor-Active Compounds in Grapes and Wines from Vitis vinifera and Non-Foxy American Grape Species. Journal of Agricultural and Food Chemistry, 59, 10657–10664.

Trott, O. and Olson, A.J. (2010) AutoDock Vina: improving the speed and accuracy of docking with a new scoring function, efficient optimization, and multithreading. J Comput Chem, 31, 455–461.

Turlings, T.C., Tumlinson, J.H. and Lewis, W.J. (1990) Exploitation of herbivore-induced plant odors by host-seeking parasitic wasps. Science, 250, 1251–1253.

von Merey, G.E., Veyrat, N., D’Alessandro, M. and Turlings, T.C. (2013) Herbivore-induced maize leaf volatiles affect attraction and feeding behavior of Spodoptera littoralis caterpillars. Front Plant Sci, 4, 209.

Wang, J. and De Luca, V. (2005) The biosynthesis and regulation of biosynthesis of Concord grape fruit esters, including ‘foxy’ methylanthranilate. Plant J, 44, 606–619.

Westfall, C.S., Muehler, A.M. and Jez, J.M. (2013) Enzyme action in the regulation of plant hormone responses. J Biol Chem, 288, 19304–19311.

Yadav, G.D. and Krishnan, M.S. (1998) An Ecofriendly Catalytic Route for the Preparation of Perfumery Grade Methyl Anthranilate from Anthranilic Acid and Methanol. Organic Process Research & Development, 2, 86–95.

Yang, Y., Cuenca, J., Wang, N., Liang, Z., Sun, H., Gutierrez, B., Xi, X., Arro, J., Wang, Y., Fan, P., Londo, J., Cousins, P., Li, S., Fei, Z. and Zhong, G.Y. (2020) A key ‘foxy’ aroma gene is regulated by homology-induced promoter indels in the iconic juice grape ‘Concord’. Hortic Res, 7, 67.

Yang, Z. (1994) Maximum likelihood phylogenetic estimation from DNA sequences with variable rates over sites: Approximate methods. Journal of Molecular Evolution, 39, 306–314.

Zubieta, C., Ross, J.R., Koscheski, P., Yang, Y., Pichersky, E. and Noel, J.P. (2003) Structural Basis for Substrate Recognition in the Salicylic Acid Carboxyl Methyltransferase Family. The Plant Cell, 15, 1704–1716.

