## Supporting Information for "Molecular basis of one-step methyl anthranilate biosynthesis in grapes, sweet orange, and maize"

**This file contains:**

Figures S1-16
Table S1
Data S1

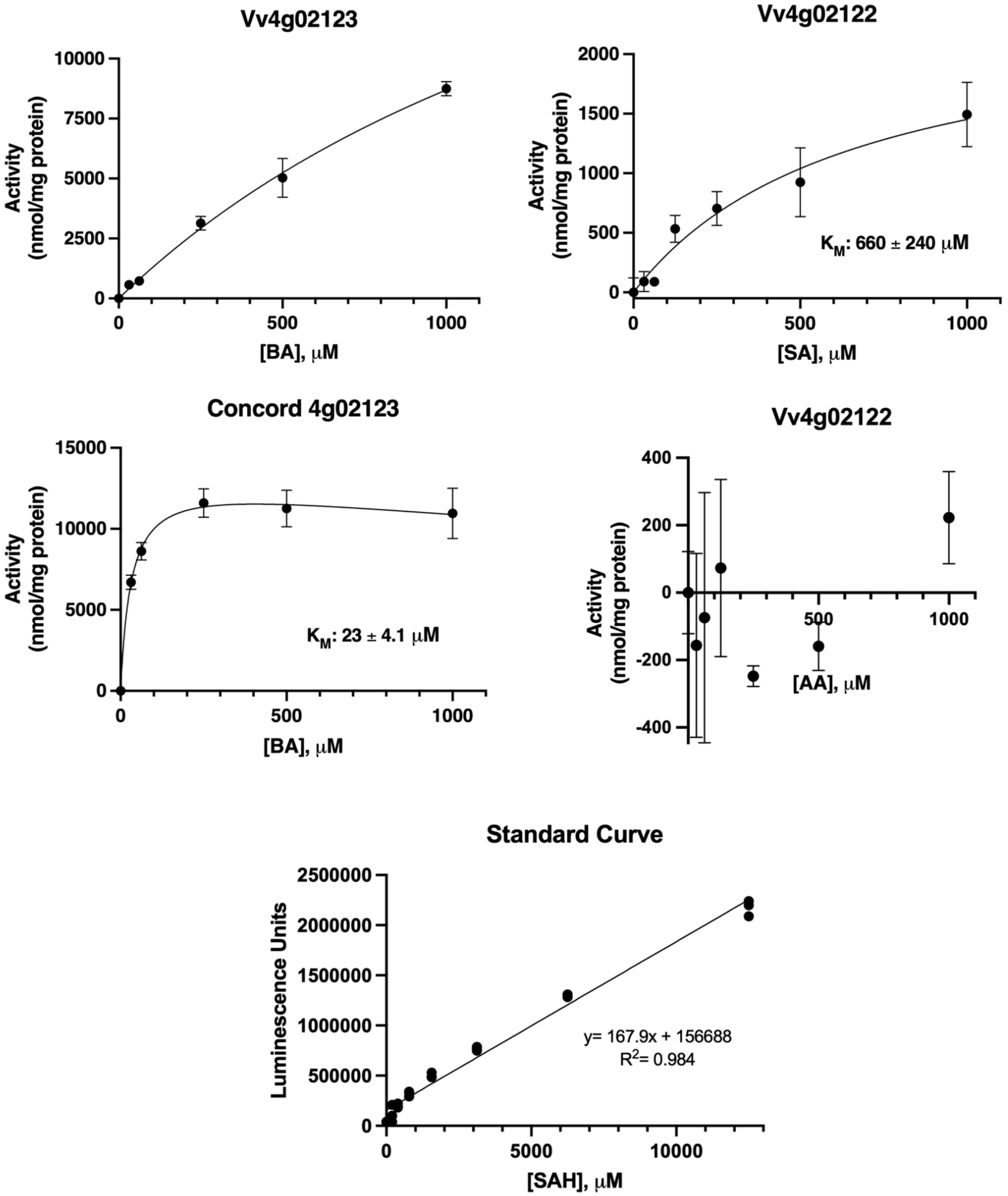


**Figure S1.** Kinetic plots of the activities of *Vitis vinifera* (Vv) 4g02122, 4g02123, and 'Concord' ortholog of 4g02123 with BA or SA. Error bars represent standard deviations; where error bars are not visible, standard deviation is too small to visualize. Standard errors are included for Km values. An SAH standard curve was used to convert luminescence units to activities with the MTase-Glo coupled assay mix. For all points, n=3.

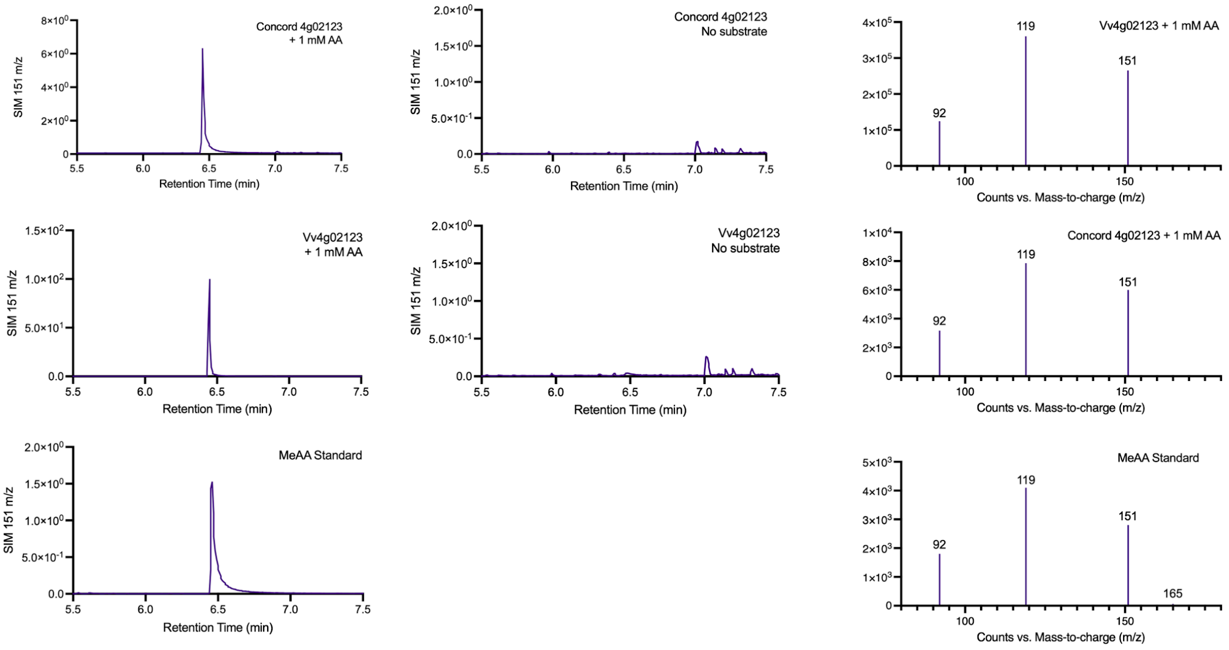


**Figure S2.** GC-MS data confirms that 4g02123 from ‘Concord’ and wine grapes (Vv) synthesizes methyl anthranilate (MeAA) *in vitro*. Representative GC-MS chromatograms from triplicate replicates are shown for selected-ion monitoring (SIM) for 151 m/z. Spectra for methyl anthranilate are shown for the 6.45-minute retention time.

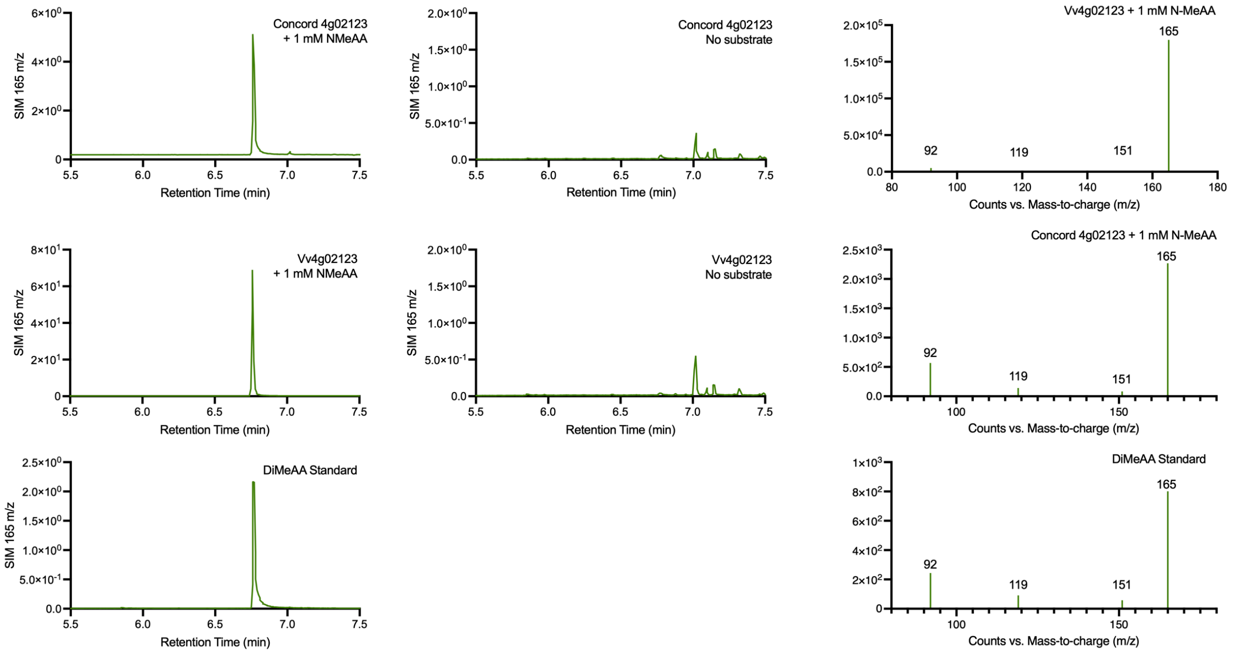


**Figure S3.** GC-MS data confirms that 4g02123 from ‘Concord’ and wine grapes (Vv) synthesizes dimethyl anthranilate (DiMeAA) *in vitro*. Representative GC-MS chromatograms from triplicate replicates are shown for selected-ion monitoring (SIM) for 165 m/z. Spectra for DiMeAA are shown for the 6.76-minute retention time. Each chromatogram is representative of triplicate reactions.

```
          1                                                                                                                    120
vinifera                                                                                                      TATTGTAATTTA--ACTTTA
labrusca  CGTTGTGATAGAAGTTAGATCGGTTTGGTTCAACTGTGCTAAAATCTTTTCAGTATTTGGCCATCTTGCCTTTTTTTTTTTTTTCATCCCTCGGCATTATTATTGGGAAGAAGTACTAGG
 Concord    TTGTGATAGAAGTTAGATCGGTTTGGTTCAACTGTGCTAAAATCTTTTCAGTATTTGGCCATCTTGCCTTTTTTTTTTTTT-CATCCCTTGGCATTATTATTGGGAAGAAGTACTAGG

          121                                                                                                                  240
vinifera  AAAACCTTAAAT-TCAAAAAATGTAAAATTTGGAGAGTACGGAAAAAAATTAGGACGATCTAATTACCTAA--------GCTTTAC-ATATGATTATCAAAATGTTGAATTTTTATGAAA
labrusca  CCATCTTTCTGTCTCTTCAATTACTAATGGAAATTAATTCCCAAAAAGAAGTGTTCGGCCATGTATCCAAACAGTGCTAGCTGGCCGATATTTTTATCAGAATATAAGGTCCTCATTCGT
 Concord  CCATCTTTCTGTCTCTTCAATTACTAATGGAAATTAATTCCCAAAAAGAAGTGTTCGGCCATGTATCCAAACAGTGCTAGCTGGCCGATATTTTTATCAGAATATAAGGTCCTCATTCGT

          241                                                                                                                  360
vinifera  ATACTCATTTCATATTAAAATATTACCATATT-CATTTTTATGTTGAATTTAAATGTTTAATAATTTGTTAGAAATAAAATAATGTAATTATGATATATGATGATTTTTTTTGTTAATAA
labrusca  TTCCACAAACCATTATGGAAAG--ACTGTGTGGCATCTTCATGCCAGAATAGAATGCACTCGCTGCACTCAGACACTTGTTTATGTGGCTCAGATTATGGAAGATATCACATGGCCTAAG
 Concord  TTCCACAAACCATTATGGAAAG--ACTGTGTGGCATCTTCATGCCAGAATAGAATGCACTCGCTGCACTCAGACACTTGTTTATGTGGCTCAGATTATGGAAGATATCACATGGCCTAAG

          361                                                                                                                  480
vinifera  ATATGTTTTCAAATTTTATATTATTTTTATTTTTATTTTTTT--TCCA--ATTCTAT---TTTTATCTAATATTTATTTTATTTAAACC--CAAAATTGTTGAGAATGGAGGAAGAAAGA
labrusca  AAATGTGTAGGGTTGCTAAGAAAGCGTCATGCTCCGCTGAGACGTCCAGTATTCTGTGGCTGATGTCTAACCCCAATGTGTTTAAGCAGGCTATCCTGGCCCAAGCGGAAAACTCATGC
 Concord  AAATGTGTAGGGTTGCTAAGAAAGCGTCATGCTCCGCTGAGACGTCCAGTATTCTGTGGCTGATGTCTAACCCCAATGTGTTTAAGCAGGCTATCCTGGCCCAAGCGGAAAACTCATGC

          481                                                                                                                  600
vinifera  ATGCCACATCCCGTT------CTTCCCTTAACTG-GTTAAGTGGAATTTTTATTT-CAAATTTAAAATGGATGC-CCAAGATGT------AATCTTGGACATCCAATAGCTTGAAAAACT
labrusca  ACCCACCTTCACTTTGTCCTACTTTTCATCATTGTGTTTCATGCACGCATTATTAGCATTATTTTTGTTGTTTCTCCGTTGTTTTTAGTCAGTATTGGTGAATATAAGGAATAATACCCA
 Concord  ACCCACCTTCACTTTGTCCTACTTTTCATCATTGTGTTTCATGCACGCATTATTAGCATTATTTTTGTTGTTTCTCCGTTGTTTTTAGTCAGTATTGGTGAATATAAGGAATAATACCCA

          601                                                                                                                  720
vinifera  TTTTCGGTAGGTACCGAAGAGGTACCCTAGAATTTTCCAA--------TTCAT----------TTCACACCTAAACTGAAGATTCAAACTACAAACATTCATCTTGTCATTGAAAATTAT
labrusca  TCAGTGGTGTCAGTTGAAGAACTTTCCTTGTCACGGCCTAAAACTTACTTCAAGGGCATGAAATTCATTTCACACCTGAAGATTCGAAGTACAAACATTCATCTTGTAATTGAAAATTAT
 Concord  TCAGTGGTGTCAGTTGAAGAACTTTCCTTGTCACGGCCTAAAACTTACTTCAAGGGCATGAAATTCATTTCACACCTGAAGATTCGAAGTACAAACATTCATCTTGTAATTGAAAATTAT

          721                                                                                                                  840
vinifera  TAATTACTAATTATCAAATTTATGTTCAAATAAAAATTCTAAAATATTCTATTCTCAACTTTTAAACCAACATATCAATCCAAGTGTTATTGTCCAAATAACT-CTAAATTCAAATAATTC
labrusca  TAATTACTAATTATCAAATTTATGTTCAGATAAAAATTCTAAAATCTTCTATTCTCAACTTTTAAACCAACATATCAACCCAAGTATTATTGTCCAAATAACTTCAAAATTCAAATAATTC
 Concord  TAATTACTAATTATCAAATTTATGTTCAGATAAAAATTCTAAAATCTTCTATTCTCAACTTTTAAACCAACATATCAACCCAAGTATTATTGTCCAAATAACTTCAAAATTCAAATAATTC

          841                                                                                                                  960
vinifera  GAATGGAAATTAAATTTTAACATCTAAAATAAAATTTGAACTAAAATCCTAATAAAGAAAAATTCTCCAACCTAACCATCACTCCTCATTCGATCTGAGAGTATCTGAAAAATTATCAATA
labrusca  AAACGGAAATTAAATTTTAACATCCAAAATAAAATTTGAACCAAAATCCTAATAAAGAAAAATTCTCCAACCTAACCATCACTCCTCATTCGAACTGAGAGTATCTGAAAAATTATCAACA
 Concord  GAACGGAAATTAAATTTTAACATCCAAAATAAAATTTGAACCAAAATCCTAATAAAGAAAAATTCTCCAACCTAACCATCACTCCTCATTCGAACTGAGAGTATTTGAAAAATTATCAACA

          961                                                                                                                 1080
vinifera  AAGAGGCATGAACTTAAAACTTAATAAAGAACATCAATACAATTTCATCGATAAAACATTTTCAATTACAATTACAAATCAGAGATACAATATATCATTATTTTCATAAAAACTTTTCAGT
labrusca  AAAAGGCATGAACTTAAAACTTAGTAAGGAACATCAATACAATTTTATAGATCAAACATTTTTAATTACAATTACAAATGGGAGATACAATATATCATTATTTTCACAAAAGCTTTTTAGT
 Concord  AAAAGGCATGAACTTAAAACTTAGTAAGGAACATCAATACAATTTTATAGATCAAACATTTTTAATTACAATTACAAATGGGAGATACAATATATCATTATTTTCACAAAAGCTTTTTAGT

          1081                                                                                                                1200
vinifera  AAAACTTTCTCATATCTATTCCAGAATAATTTCTCATCAAAAGGAAATTAAATACATTCAAAATATCTTTACTCTGGTTATTAAATAACAAATAGTGCCTAATTAGATGAGACTTCACAAT
labrusca  AAAATTTTCTCATATCTATTCCAAAATAATTTCTCATCAAAAGCAAATTAAATACATTCAAAATATTTTTACTCTGATTATCGAATAACAAATAGTGCCTAATTAGGTGAGACTTCACAAT
 Concord  AAAATTTTCTCATATCTATTCCAAAATAATTTCTCATCAAAAGCAAATTAAATACATTCAAAATATTTTTACTCTGACTATAGAATAACAAATAGTGCCTAATTAGGTGAGACTTCACAAT

          1201                                                                                                                1320
vinifera  ATATTATTATTATTATTTCTTTCAATTACCAGTAACACTATTACTCCAAGTCACCATATATTATTATTATTATTTCTGATGGTTGTATATTATTATTTTTTTTTGTCACTCAATAATCCGC
labrusca  ATATTATTATTATTATT-------------------------------------------------------------------------------------------------------
 Concord  ATATTATTATTATTATT-------------------------------------------------------------------------------------------------------

          1321                                                                                                                1440
vinifera  TCGTATTTGTTCTTCTTTTCTTTATTATTATTATTATTATTATTATTATTATTATTATTATTATTATTTTCT---CAAATTGTCGCTACATAGTTCCTCAAATTCTAAAATCCAGAAA
labrusca  ------------------------------------------------------------ATTATTATTTTCT---CAAATTGTCGCTACATAGTTCCTAAAATTCTAAAATCCAGAAA
 Concord  ------------------------------------------------------------ATTATTATTATTTTCTCAAATTGTCGCTACATAGTTCCTAAAATTCTAAAATCCAGAAA

          1441                                                                                                                1560
vinifera  CATGCAATCATTTTTC-----------------------TTATTATTTAGTTTTTCCGATCTAAAGCCTCCACAAACAATTTATTGATTTATTTTCTAAATCTAAAATTTCTCATCTTCCA
labrusca  CATGCAATCATTTTTCTTATTATTAATTTAATTTAATTTTTATTATTTAGTTTTTCCGATCTAAAGCCTCCACAAACAATTTATTGATTTATTTTCTAAATCTAAAATTTCTCATCTTCCA
 Concord  CATGCAATCATTTTTCTTATTATTAATTTAATTTAATTTTTATTATTTAGTTTTTCCGATCTAAAGCCTCCACAAACAATTTATTGATTTATTTTCTAAATCTAAAATTTCTCATCTTCCA

          1561                                                                                                                1680
vinifera  CTATTTATTTATTTATTTTCTATTCAAGTGAAATTTTCCTATTTTCATTAATAAATTAATGTACATTAATAAATTTCCAATCTTTTGATCTAAAAAATTAAATAAACAAACCCTAATTTAG
labrusca  CTATTTATTTATTTATTTTCTATTCAATTGAAATTTTCCTATTTTCATCAATAAATTAATCTACATTAATAAATTTCCAATTTTTTGATCTAAAAAATTAAATAAACAAACCCTAATTTAG
 Concord  CTATTTATTTATTTATTTTCTATTCAATTGAAATTTTCCTATTTTCATCAATAAATTAATCTACATTAATAAATTTCCAATTTTTTGATCTAAAAAATTAAATAAACAAACCCTAATTTAG

          1681                                                                                                                1800
vinifera  ATTGTGATTTTGAAAGTATTCAGATCTAATTAACGAATATAAAATACTAATTCGAGGATTAAAATCTTACCTAAAAATTTGATTTCAAACCCTAAACCTCCAAATCTTTAACCCTATACT
labrusca  ATTGTAATTTCTAAAGTATTCAAATCTAATTAACGAATATAAAATACTAATTCGAGGATTAAAATCTTACCTAAAAATTTGATTTCAAACCCTAAACCTCCAAATCTTCAACCCTATACT
 Concord  ATTGTAATTTCTAAAGTATTCAAATCTAATTAACGAATATAAAATACTAATTCGAGGATTAAAATCTTACCTAAAAATTTGATTTCAAACCCTAAACCTCCAAATCTTCAACCCTATACT

          1801                                                                                                                1920
vinifera  CTTTGGTATTTACGGTCTTCATGAAAAAGGGTTTTTTTGAATAAAAAGATAGGAAAAAAGTAATTTTATAACTGAAGCTTTTAAAAACGAAAAGACTTTTTTTACCGTTATTAAATTAAT
labrusca  CTATGGTATTTATGGTCTTCGTGAAAAAGGGTTTTT--GAATAGAAAAGATAGTAAAAAATTAATTTTATACCTGAGGCTTTTAAAAACGAAAAGACTTTTTT-ACTGTTATTAAATTAAT
 Concord  CTATGGTATTTATGGTCTTCGTGAAAAAGGGTTTTT--GAATAGAAAAGATAGTAAAAAATTAATTTTATACCTGAGGCTTTTAAAAACGAAAAGACTTTTTT-ACTGTTATTAAATTAAT

          1921                                                                                                                2040
vinifera  TTAATTAATTAATTAATTAATTTCTAAATTATAAATTTACCCCTAAATTGGGTTTGGGGGTATTACATTCCTTTTGAATTATCACCAACAATGACCATCGGGGGGGTTACGTGAAAGTAGA
labrusca  TTAATTAATTAATTAATT----TCTAAATTATAAATTTACCCCTAAATTGGGTT-GGGGGTATTACATTCCTTTTGAATTATCACCAACAGTGACCAGCGGGG---TTACGTGGAAGTAGA
 Concord  TTAATTAATTAATTAATT----TCTAAATTATAAATTTACCCCTAAATTGGGTT-GGGGGTATTACATTCCTTTTGAATTATCACCAACAGTGACCAGCGGGG---TTACGTGGAAGTAGA

          2041                                                                                                                2160
vinifera  GTACATAGACGATGAAATAATTGATTCCACAACGATGTGATTCCAAGTTCTATAAATAGTGAGCATCTCCTTCAACACAATTCACATTGCAACTGAGAGAGAGCAATCAGCAAAGAGGCAG
labrusca  GTACATAGACGATGAAATAATTGATTCCACAACGATGTGACTCCAAGTTCTATAAATAGTGAGCATCTCCTTCAACACAATTCACATTGCAACTGAGAGAGAGCAATCAGCAAAGAGGCAG
 Concord  GTACATAGACGATGAAATAATTGATTCCACAACGATGTGACTCCAAGTTCTATAAATAGTGAGCATCTCCTTCAACACAATTCACATTGCAACTGAGAGAGAGCAATCAGCAAAGAGGCAG

           2161  2183
vinifera  GAAGGGAGAG(ATG)
labrusca  GAAGGGAGAG(ATG)
 Concord  GAAGGGAGAG(ATG)
```

**Figure S4**. Promoter sequence comparisons for *Vitis vinifera* Vv4g02123 and its orthologs in *Vitis labrusca* and ‘Concord’ grapes. The start codon is highlighted in green.

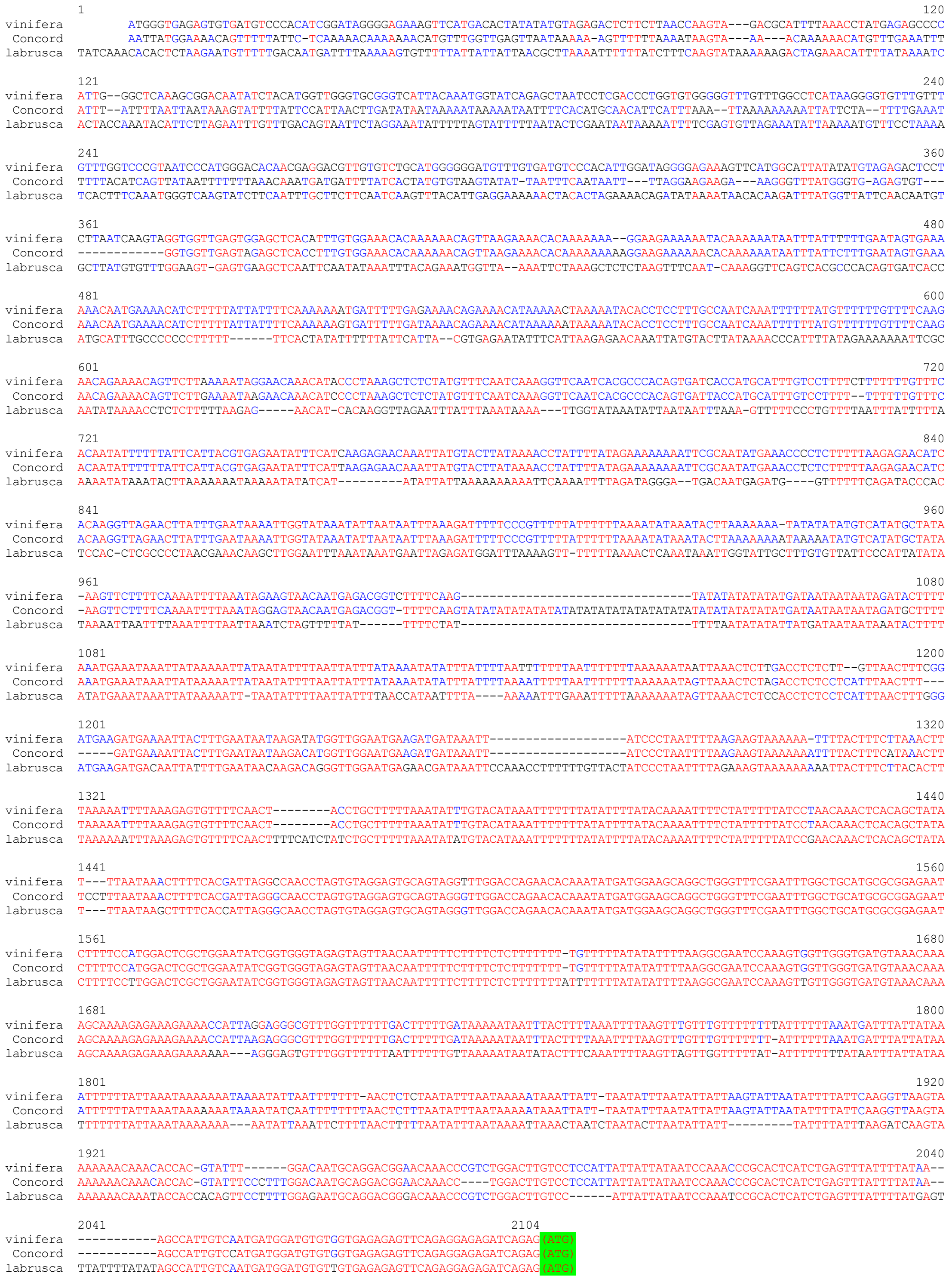

**Figure S5.** Promoter sequence comparisons for *Vitis vinifera* Vv4g02169 and its orthologs in *Vitis labrusca* and 'Concord' grapes. The start codon is highlighted in green.

```
           1                                                                                                  100
 vinifera  ATGGAAGTAGTTCAAGTGCTTTGCATGAAGGGAGGAAATGGGGATACCAGTTACGCAAAAAACTCATTAGTTCAGAAAAAAGTAATATCCTTGACAAAGC
 labrusca  ATGGAAGTAGTTCAAGTGCTTTGCATGAAGGGAGGAAATGGGGATACCAGTTACGCAAAAAACTCATTAGTTCAGAAAAAAGTAATATCCTTGACAAAGC
  Concord  ATGGAAGTAGTTCAAGTGCTTTGCATGAAGGGAGGAAATGGGGATACCAGTTACGCAAAAAACTCATTAGTTCAGAAAAAAGTAATATCCTTGACAAAGC

           101                                                                                                200
 vinifera  CCATAATCGAGGAAGCCATAACAAATCTTTACTGCAACAAATTTCCGACCAGCCTATGCATCGCAGACTTGGGATGTTCTTCTGGACCCAACACTTTGTT
 labrusca  CCATAATCGAGGAAGCCATAACAAATCTTTACTACAACAAATTTCTGACCAGCCTATGCATCGCAGACTTGGGATGTTCTTCTGGACCCAACACTTTGTT
  Concord  CCATAATCGAGGAAGCCATAACAAATCTTTACTACAACAAATTTCTGACCAGCCTATGCATCGCAGACTTGGGATGTTCTTCTGGACCCAACACTTTGTT

           201                                                                                                300
 vinifera  TGCGGTCTTGGAAGTTGTCACTACAGTGGACAGGGTGGGCAAGAAAATGGGGCGTCAATTGCCCGAAATTCAAGTGTTTTTGAATGATCTGCCGGGGAAT
 labrusca  TGCGGTCTTGGAAGTTGTCACTACAGTGGACAGGGTGGGCAAGAAAATGGGGCGTCAATTGCCCGAAATTCAAGTGTTTTTGAATGATCTGCCGGGGAAT
  Concord  TGCGGTCTTGGAAGTTGTCACTACAGTGGACAGGGTGGGCAAGAAAATGGGGCGTCAATTGCCCGAAATTCAAGTGTTTTTGAATGATCTGCCGGGGAAT

           301                                                                                                400
 vinifera  GATTTCAACACCATTTTCAAATCCTTGCCCAGGTTCCAAAAGGATCTTGAGAAAAGAATGGGAGCAGGAGCTGAATCATGTTTCATAAACGGAGTTCCAG
 labrusca  GATTTCAACACCATTTTCAAATCCTTGCCCGGGTTCCAAAAGGATCTTGAGAAAAGAATGGGAGCAGGAGCTGAATCATGTTTCATAAACGGAGTTCCAG
  Concord  GATTTCAACACCATTTTCAAATCCTTGCCCGGGTTCCAAAAGGATCTTGAGAAAAGAATGGGAGCAGGAGCTGAATCATGTTTCATAAACGGAGTTCCAG

           401                                                                                                500
 vinifera  GTTCTTTCTACGGCAGGCTGTTCCCCAGCAAAAGTCTTCATTTTATCCATTCTTCTTACAGTCTCCAATGGTTGTCTCAGGTTCCTCAAGGGCTGGAGAG
 labrusca  GTTCTTTCTACGGCAGGCTGTTCCCCAGCAAAAGTCTTCATTTTATCCATTCTTCTTACAGTCTCCAATGGTTGTCTCAGGTTCCTCAAGGGCTGGAGAG
  Concord  GTTCTTTCTACGGCAGGCTGTTCCCCAGCAAAAGTCTTCATTTTATCCATTCTTCTTACAGTCTCCAATGGTTGTCTCAGGTTCCTCAAGGGCTGGAGAG

           501                                                                                                600
 vinifera  TAACAAAGGGAACATTTATATGGCAAGTTCGAGCCCACCATGCGTGCTTAAAGTATACTATGAGCAATTCCGAACCGATTTCTCCATGTTCCTCAGGTGT
 labrusca  TAACAAAGGGAACATTTATATGGCAAGTTCGAGCCCACCATGCGTGCTTAAAGTATACTATGAGCAATTCCGAACCGATTTCTCCATGTTCCTCAGGTGT
  Concord  TAACAAAGGGAACATTTATATGGCAAGTTCGAGCCCACCATGCGTGCTTAAAGTATACTATGAGCAATTCCGAACCGATTTCTCCATGTTCCTCAGGTGT

           601                                                                                                700
 vinifera  CGGTCGGAGGAACTTCTAGAAGGAGGGAGTATGGTTTTAACATTTTTAGGAAGAAGAAGTGAAGATCCCTCTAGCAAAGAATGTTGCTACATTTGGGAGC
 labrusca  CGGTCGGAGGAACTTCTAGAAGGAGGGAGTATGGTTTTAACATTTTTAGGAAGAAGAAGTGAAGATCCCTCTAGCAAAGAATGTTGCTACATTTGGGAGC
  Concord  CGGTCGGAGGAACTTCTAGAAGGAGGGAGTATGGTTTTAACATTTTTAGGAAGAAGAAGTGAAGATCCCTCTAGCAAAGAATGTTGCTACATTTGGGAGC

           701                                                                                                800
 vinifera  TCTTAGCTGTGGCTCTCAACGATATGGTGGCAGAGGGGCTCATAGAGGAGGAAAAAATGGATTCCTTCAATATTCCTCAATATACACCATCCCCAGCTGA
 labrusca  TCTTAGCTGTGGCTCTCAACGATATGGTGGCAGAGGGGCTCATAGAGGAGGAAAAAATGGATTCCTTCAATATTCCTCAATATACACCATCCCCAGCTGA
  Concord  TCTTAGCTGTGGCTCTCAACGATATGGTGGCAGAGGGGCTCATAGAGGAGGAAAAAATGGATTCCTTCAATATTCCTCAATATACACCATCCCCAGCTGA

           801                                                                                                900
 vinifera  AGTAAAATGTGAGGTTGAAAAGGAAGGATCCTTTACCATAAGTAAGCTGGAGGTTTCTGAAGTTAACTGGAACGCGTATCACGGTGAATTCTGCCCATCA
 labrusca  AGTAAAATGTGAGGTTGAAAAGGAAGGATCCTTTACCATAAGTAGGCTGGAGGTTTCTGAAGTTAACTGGAACGCGTATCACGGTGAATTCTGCCCATCA
  Concord  AGTAAAATGTGAGGTTGAAAAGGAAGGATCCTTTACCATAAGTAGGCTGGAGGTTTCTGAAGTTAACTGGAACGCGTATCACGGTGAATTCTGCCCATCA

           901                                                                                               1000
 vinifera  GATGCTCATAAAGATGGTGGGTACAATGTGGCTAAGTTGATGAGAGCAGTGGCGGAGCCATTGCTTGTAAGCCATTTCGGTGATGGAATAATAGAGGAAG
 labrusca  GATGCTCATAAAGATGGTGGGTATAATGTGGCTAAGTTGATGAGAGCAGTGGCGGAGCCATTGCTTGTAAGCCATTTCGGTGATGGAATAATAGAGGAAG
  Concord  GATGCTCATAAAGATGGTGGGTATAATGTGGCTAAGTTGATGAGAGCAGTGGCGGAGCCATTGCTTGTAAGCCATTTCGGTGATGGAATAATAGAGGAAG

           1001                                                                                 1095
 vinifera  TGTTCAGCAGGTACCAAAAGATTGTGGCTGATCGCATGTCCAGAGAGAAGACTGAGTTCGTAAATGTCACTGTCTCCATGACTAAGCGTGGATAA
 labrusca  TGTTCAGCAGGTACCAAAAGATTGTGGCTGATCGCATGTCCAGAGAGAAGACTGAGTTCGTAAATGTCACTGTCTCCATGACTAAGCGTGGATAA
  Concord  TGTTCAGCAGGTACCAAAAGATTGTGGCTGATCGCATGTCCAGAGAGAAGACTGAGTTCGTAAATGTCACTGTCTCCATGACTAAGCGTGGATAA
```

**Figure S6.** Coding sequence alignment of *Vitis vinifera* Vv4g02123 and its orthologs in *Vitis labrusca* and 'Concord' grapes.

```
           1                                                                                                   100
vinifera   ATGAAGAAGAAGGAGAGCATGGGAGTGCAGCAAGTTATCTGCATGAAAGGAGGTGTCGGAGAGGGAAGCTATGCCCGTAACTCCAAATCACAGGCTGCAT
 Concord   ATGAAGAAGAAGGAGAGCATGGGAGTGCAGCAAGTTATCTGCATGAAAGGAGGTGTCGGAGAGGGAAGCTATGCCCGTAACTCCAAATCACAGGCTGCAT
labrusca   ATGAAGAA---GGAGAGCATGGGAGTGCAGCAAGTTATCTGCATGAAAGGAGGTGTCGGAGAGGGAAGCTATGCCCGTAACTCCAAATCACAGGCTGCAT

           101                                                                                                 200
vinifera   TACTATCCAAGTCCATGCCCCTGCTGGAGCAAGCAGTGCTAGATTTATGTTGCACCACCTTACCTGAGAGTGTCGCCATAGCAGACCTTGGCTGTTCATC
 Concord   TACTATCCAAGTCCATGCCCCTGCTGGAGCAAGCAGTGCTAGATTTATGTTGCACCACCTTACCTGAGAGTGTCGCCATAGCAGACCTTGGCTGTTCATC
labrusca   TACTATCCAAGTCCATGCCCCTGCTGGAGCAAGCAGTGCTAGATTTATGTTGCACCACCTTACCTGAGAGTGTCGCCATAGCAGACCTTGGCTGTTCATC

           201                                                                                                 300
vinifera   GGGCCCAAATACTTTTTTCGCGGTCTCCGAAATCATGACCATCATCTACAGGAGGTGTCGCCAACTAGGCCGGTCACCACCAGGTTTTTGGGTGTTCTTG
 Concord   GGGCCCAAATACTTTTTTCGCGGTCTCCGAAATCATGACCATCATCTACAGGAGGTGTCGCCAACTAGGCCGGTCACCACCAGGTTTTTGGGTGTTCTTG
labrusca   GGGCCCAAATACTTTTTTCGCGGTCTCCGAAATCATGACCATCATCTACAGGAGGTGTCGCCAACTAGGCCGGTCACCACCAGGTTTTTGGGTGTTCTTG

           301                                                                                                 400
vinifera   AATGATCTTCCAGGGAATGACTTCAATGCTGTGTTCAAGTCATTGCCAACATTCCATGAAAAGATGAAGGAAGAAAATGGACAGGAGTTTGGGCCATGTC
 Concord   AATGATCTTCCAGGGAATGACTTCAATGCTGTGTTCAAGTCATTGCCAACATTCCATGAAAAGATGAAGGAAGAAAATGGACAGGAGTTTGGGCCATGTC
labrusca   AATGATCTTCCAGGGAATGACTTCAATGCTGTGTTCAAGTCATTGCCAACATTCCATGAAAAGATGAAGGAAGAAAATGGACAGGAGTTTGGGCCATGTC

           401                                                                                                 500
vinifera   ATGTTGCTGCTGTTCCGGGTTCTTTCTACCACAAGCTTTTTCCATCCAGGAGACTACACTTTGTGCACTCTTCTTGCAGTCTCCATTGGCTGTCCCAGGT
 Concord   ATGTTGCTGCTGTTCCGGGTTCTTTCTACCACAAGCTTTTTCCATCCAGGAGACTACACTTTGTGCACTCTTCTTGCAGTCTCCATTGGCTGTCCCAGGT
labrusca   ATGTTGCTGCTGTTCCGGGTTCTTTCTACCACAAGCTTTTTCCATCCAGGAGACTACACTTTGTGCACTCTTCTTGCAGTCTCCATTGGCTGTCCCAGGT

           501                                                                                                 600
vinifera   TCCTCCAGAGCTTCTTAACAAGCAAATTACAAACAAGGGGAAGATTTATCTTTCAAAAACAAGCTCACCGGCCCTCATAGATGCTTACGCATCTCAGTTC
 Concord   TCCTCCAGAGCTTCTTAACAAGCAAATTTCAAACAAGGGGAAGATTTATCTTTCAAAAACAAGCTCACCGGCCCTCATAGATGCTTACGCGTCTCAGTTC
labrusca   TCCTCCAGAGCTTCTTAACAAGCAAATTTCAAACAAGGGGAAGATTTATCTTTCAAAAACAAGCTCACCGGCCCTCATAGATGCTTACGCGTCTCAGTTC

           601                                                                                                 700
vinifera   CAGAGGGATTTCTCTTTGTTTCTTAAGTTGAGATCAGAAGAAACAGTCCCCGGAGGACGCATGGTTTTGTCGCTCATGGCCAGACGAACCCCAGACCCTG
 Concord   CAGAGGGATTTCTCTTTGTTTCTTAAGTTGAGATCAGAAGAAACAGTCCCCGGAGGACGCATGGTTTTGTCGCTCATGGCCAGACGAACCCCAGACCCTG
labrusca   CAGAGGGATTTCTCTTTGTTTCTTAAGTTGAGATCAGAAGAAACAGTCCCCGGAGGACGCATGGTTTTGTCGCTCATGGCCAGACGAACCCCAGACCCTG

           701                                                                                                 800
vinifera   TCTCAGATGAGAGTTGTCTGTTGTGGGATCTATTAGCACAGGCGTTGCAGGGGTTGGTTTCAGAGGGGCTTATTGCAGAGGAGAAATTAGATTCCTACAA
 Concord   TCTCAGATGAGAGTTGTCTGTTGTGGGATCTATTAGCACAGGCGTTGCAGGGGTTGGTTTCAGAGGGGCTTATTGCAGAGGAGAAATTAGATTCCTACAA
labrusca   TCTCAGATGAGAGTTGTCTGTTGTGGGATCTATTAGCACAGGCGTTGCAGGGGTTGGTTTCAGAGGGGCTTATTGCAGAGGAGAAATTAGATTCCTACAA

           801                                                                                                 900
vinifera   TGCACCATACTATCAACCTTACACAGAGGACCTAGAAACTGGGATAGAGAATGATGGATCCTTTAGCATCAACGGCCTTGAGATCATGGTCTTACCATGG
 Concord   TGCACCATACTATCAACCTTACACAGAGGACCTAGAAACTGAGATAGAGAATGATGGATCCTTTAGCATCAACGGCCTTGAGATCATGGTCTTACCATGG
labrusca   TGCACCATACTATCAACCTTACACAGAGGACCTAGAAACTGAGATAGAGAATGATGGATCCTTTAGCATCAACGGCCTTGAGATCATGGTCTTACCATGG

           901                                                                                                1000
vinifera   GACAGTGCCAGTGGTGGACAAAATTATGATAGGCCAACAACCGCCCAGAAGATTGCAAAGTCGATGAAAGCAGTGCAAGAGCCGATGTTGGCAAGTCATT
 Concord   GACAGTGCCAGTGGTGGACAAAATTATGATAGGCCAACAACCGCCCAGAAGATTGCAAAGTCGATGAAAGCAGTGCAAGAGCCGATGTTGGCAAGTCATT
labrusca   GACAGTGCCAGTGGTGGACAAAATTATGATAGGCCAACAACCGCCCAGAAGATTGCAAAGTCGATGAAAGCAGTGCAAGAGCCGATGTTGGCAAGTCATT

           1001                                                                                               1100
vinifera   TCGGGGCAGAAATCATGGACCCTTTGTTTAAAAGGCTCATGGAGATCATTGCAGCAGATACAAGGGAGGTGGAGCATGTTTCTGTCCTTGTTTCCATGAC
 Concord   TCGGGGCAGAAATCATGGACCCTTTATTTAAAAGGCTCATGGAGATCATTGCAGCAGATACAAGGGAGGTGGACCATGTTTCTGTCCTTGTTTCCATGAC
labrusca   TCGGGGCAGAAATCATGGACCCTTTATTTAAAAGGCTCATGGAGATCATTGCAGCAGATACAAGGGAGGTGGACCATGTTTCTGTCCTTGTTTCCATGAC

           1101     1113
vinifera   CAGAAAGGCTTGA
 Concord   CAGAAAGGCTTGA
labrusca   CAGAAAGGCTTGA
```

**Figure S7.** Coding sequence alignment of *Vitis vinifera* Vv4g02169 and its orthologs in *Vitis labrusca* and ‘Concord’ grapes.

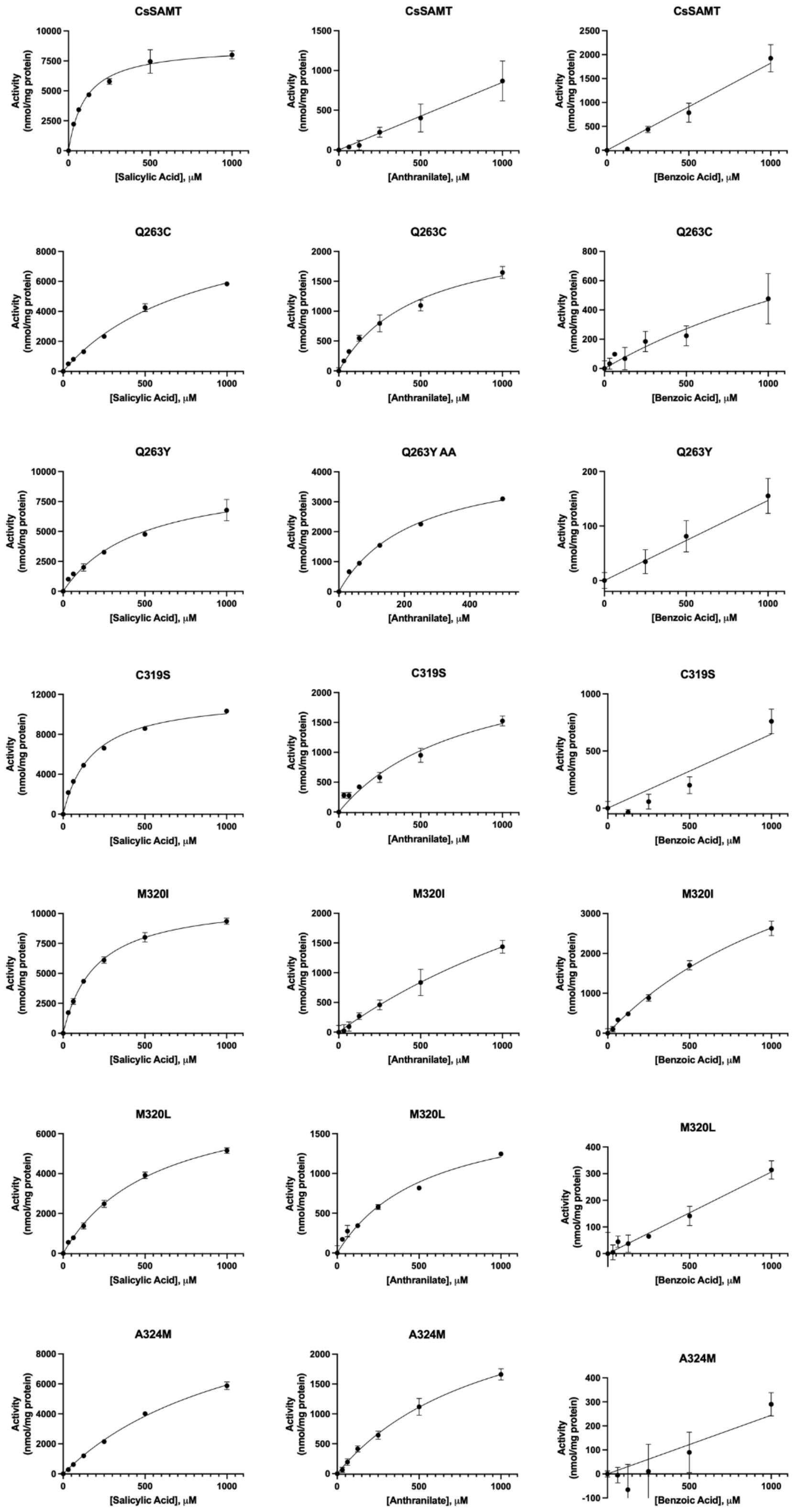


**Figure S8.** Michaelis-Menten plots of *Citrus sinensis* (Cs) SAMT activity with SA, AA or BA. Error bars represent standard deviations; where error bars are not visible, standard deviation is too small to visualize. Km values are included in Table 1. For all points, n=3.

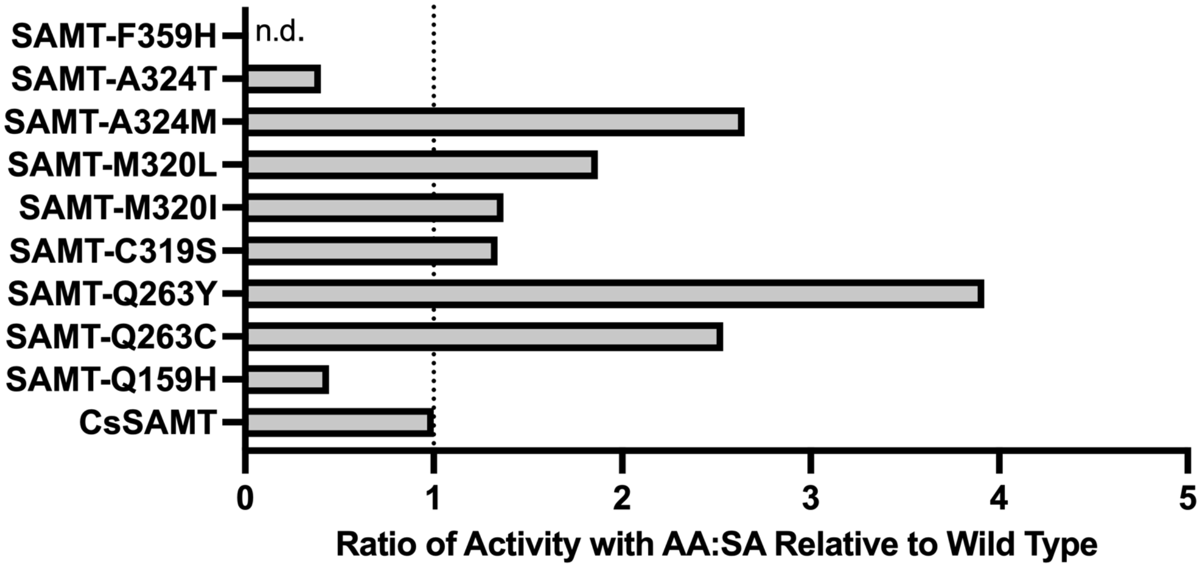


**Figure S9.** Ratio of activity with AA relative to SA for the *Citrus sinensis* SAMT. Values were calculated using 1 mM activity data in Table 1.

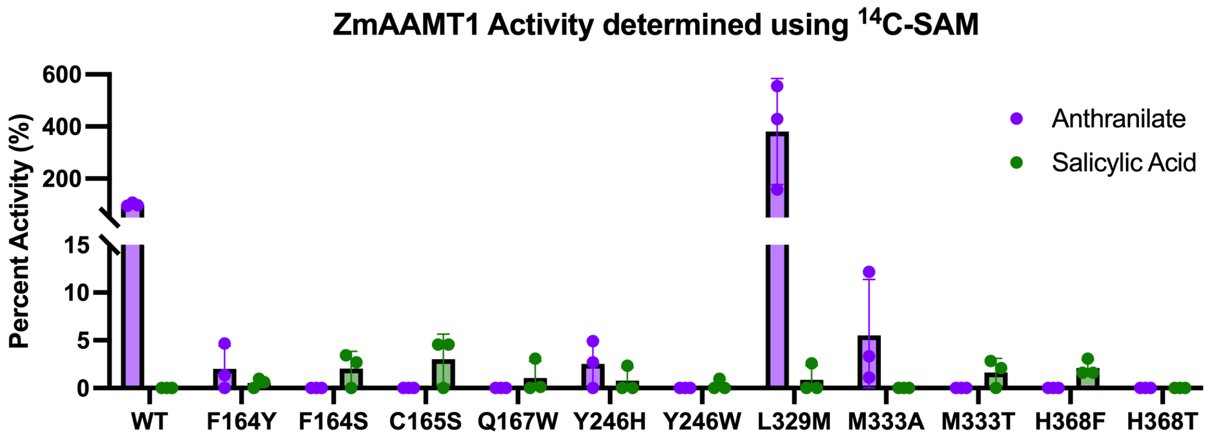


**Figure S10.** Percent activity of ZmAAMT1 mutants with 1 mM anthranilate or 1 mM salicylic acid relative to wild type, which was determined using $^{14}$C-SAM in an *in vitro* assay. Error bars represent standard deviations; where error bars are not visible, standard deviation is too small to visualize. For all points, n=3.

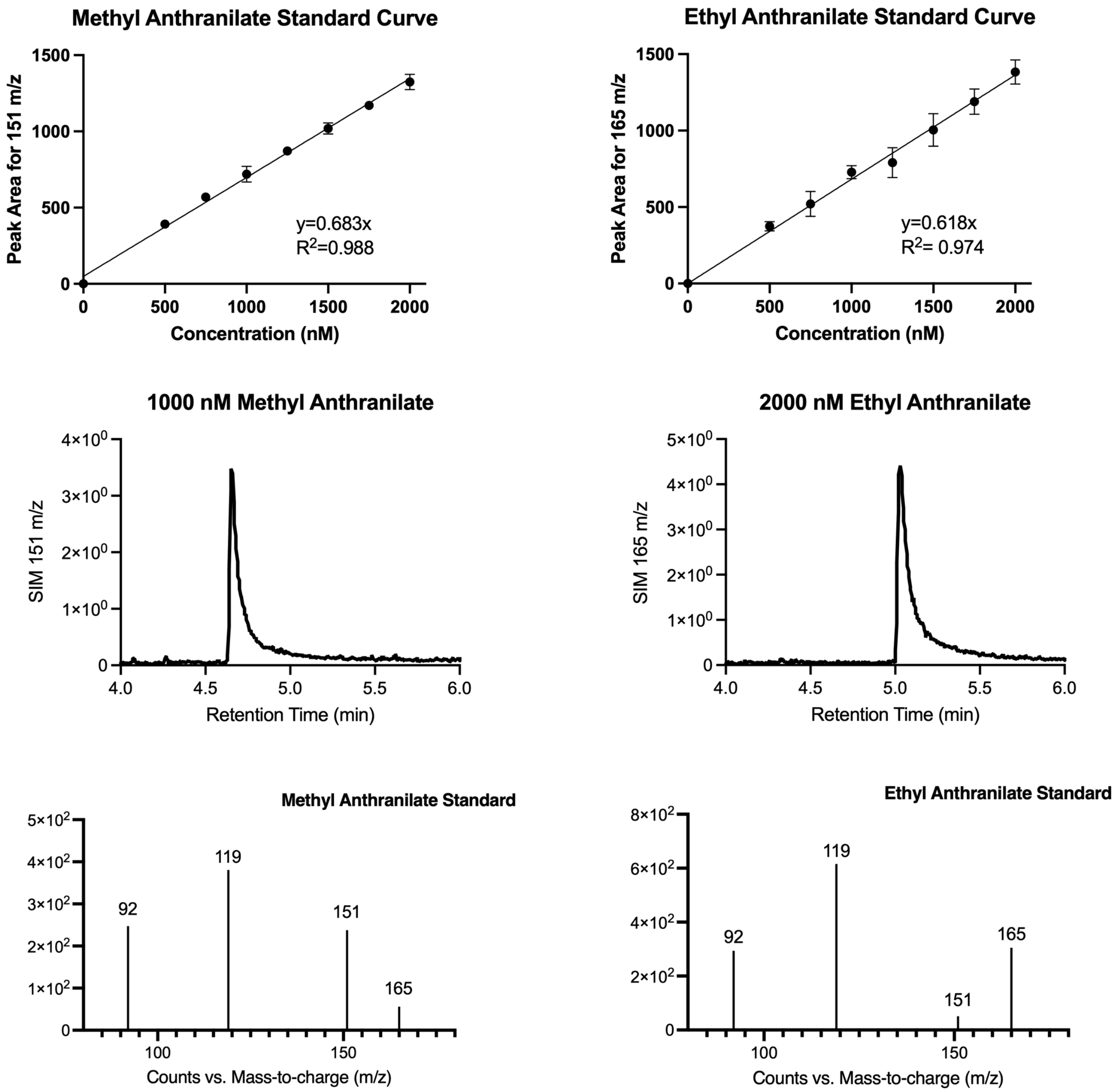


**Figure S11.** GC-MS data for *in vitro* MeAA quantification for the wild-type ZmAAMT1 enzyme and the L329M mutant. Ethyl anthranilate and methyl anthranilate calibration curves were used for the quantification of methyl anthranilate from each enzyme. Methyl anthranilate had a retention time of 4.65 minutes, while ethyl anthranilate had a retention time of 5.03 minutes. For all points, n=3.

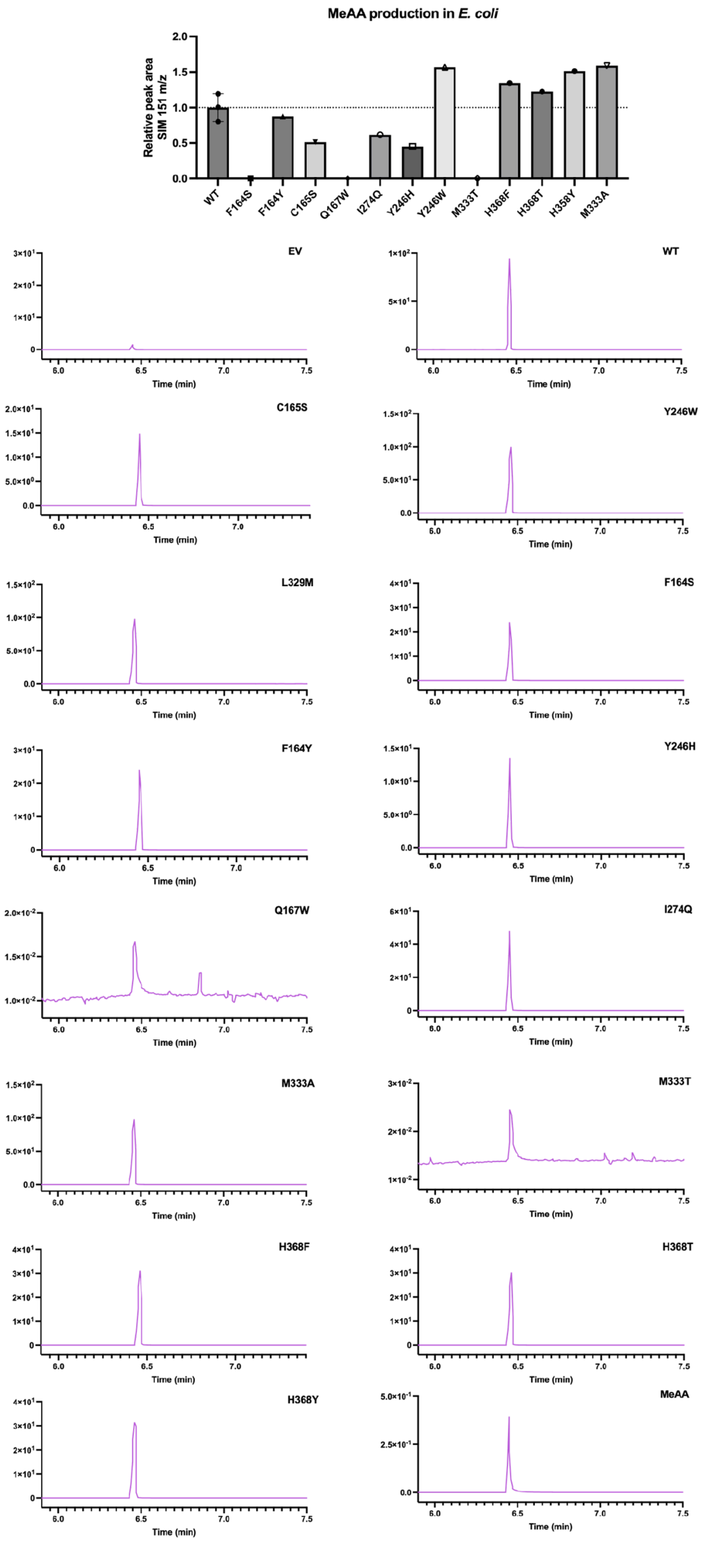


**Figure S12.** GC-MS chromatograms for *in vivo* ZmAAMT1 MeAA detection. Methyl anthranilate had a retention time of 6.45 minutes. Relative peak area was determined in comparison to *E. coli* expressing wild-type *ZmAAMT1* grown on the same day under the same conditions. Representative chromatograms for selected -ion monitoring (SIM) for 151 m/z (n=3 for WT and L329M; n=1 for all others; EV= empty vector).

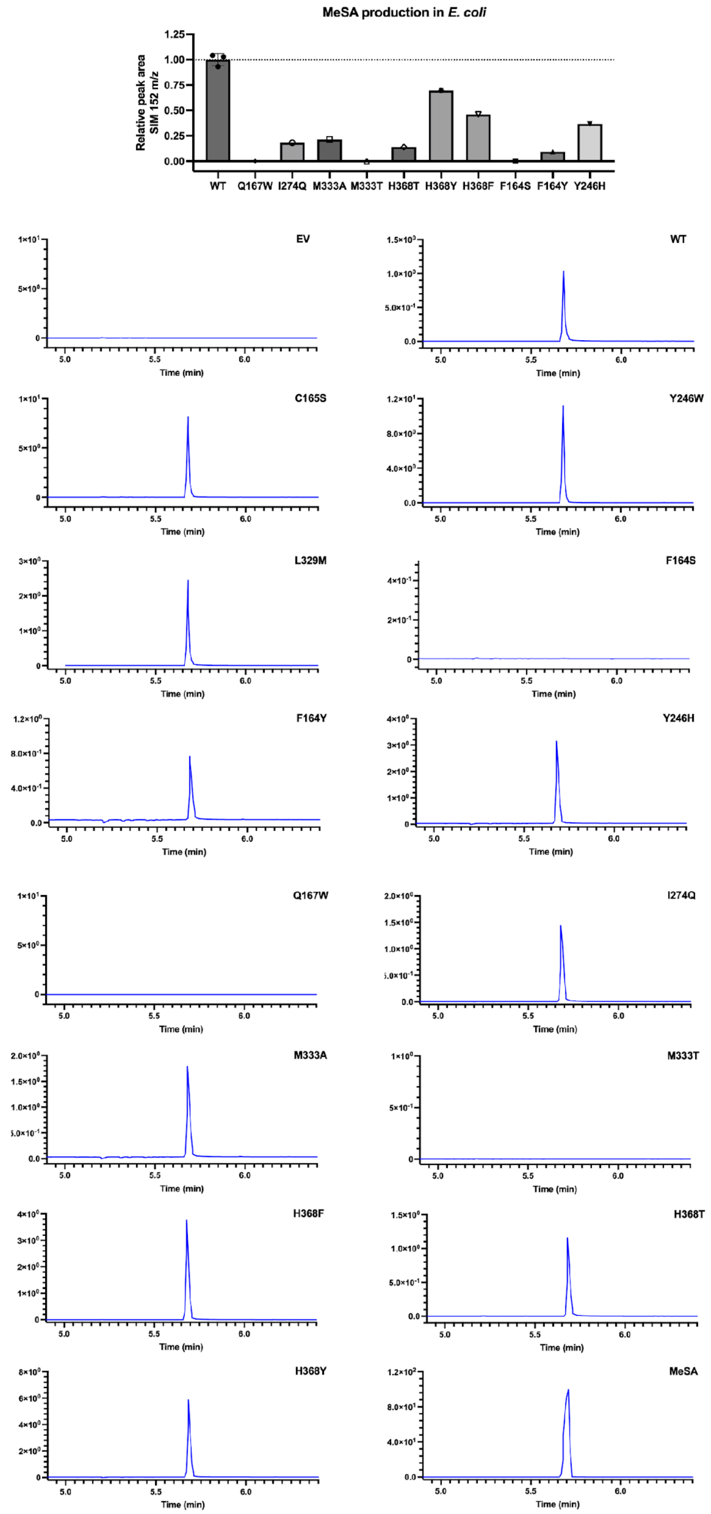


**Figure S13.** GC-MS chromatograms for *in vivo* ZmAAMT1 MeSA detection. Methyl salicylate had a retention time of 5.68 minutes. Relative peak area was determined in comparison to *E. coli* expressing wild-type *ZmAAMT1* grown on the same day under the same conditions. Representative chromatograms for selected-ion monitoring (SIM) for 152 m/z (n=3 for WT, C165S, Y246W, and L329M; n=1 for all others; EV = empty vector).

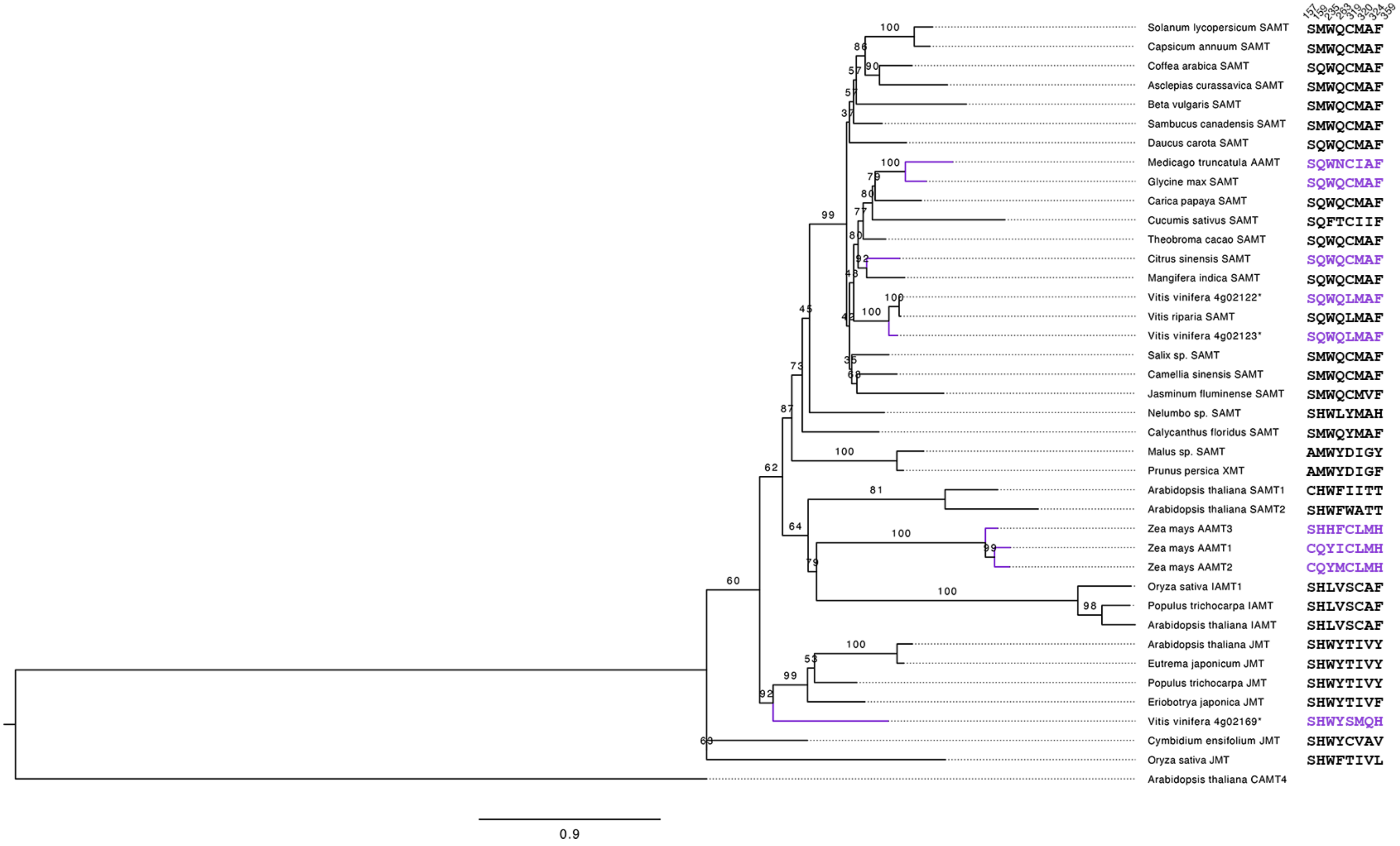


**Figure S14.** Dendrogram of plant acyl acid methyltransferases. Branches for enzymes that have been found to use anthranilate are colored purple. Asterisks are used to denote enzymes that were characterized in this study. Bootstrap values are shown as a percentage and were determined using 10,000 ultrafast bootstrap replicates. Amino acid sequence alignment on the right shows the eight active site residues that were investigated in this study. Active site amino acid numbering corresponds to the *Citrus sinensis* SAMT.

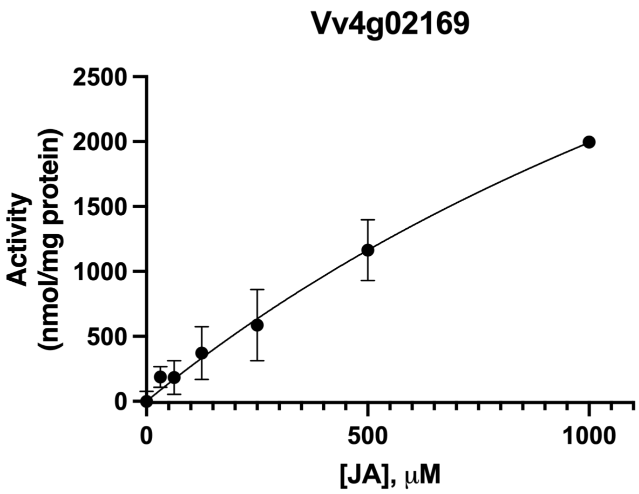


**Figure S15.** Jasmonic acid (JA) activity in the Vv4g02169 enzyme. Error bars represent standard deviations; where error bars are not visible, standard deviation is too small to visualize. For all points, n=3.

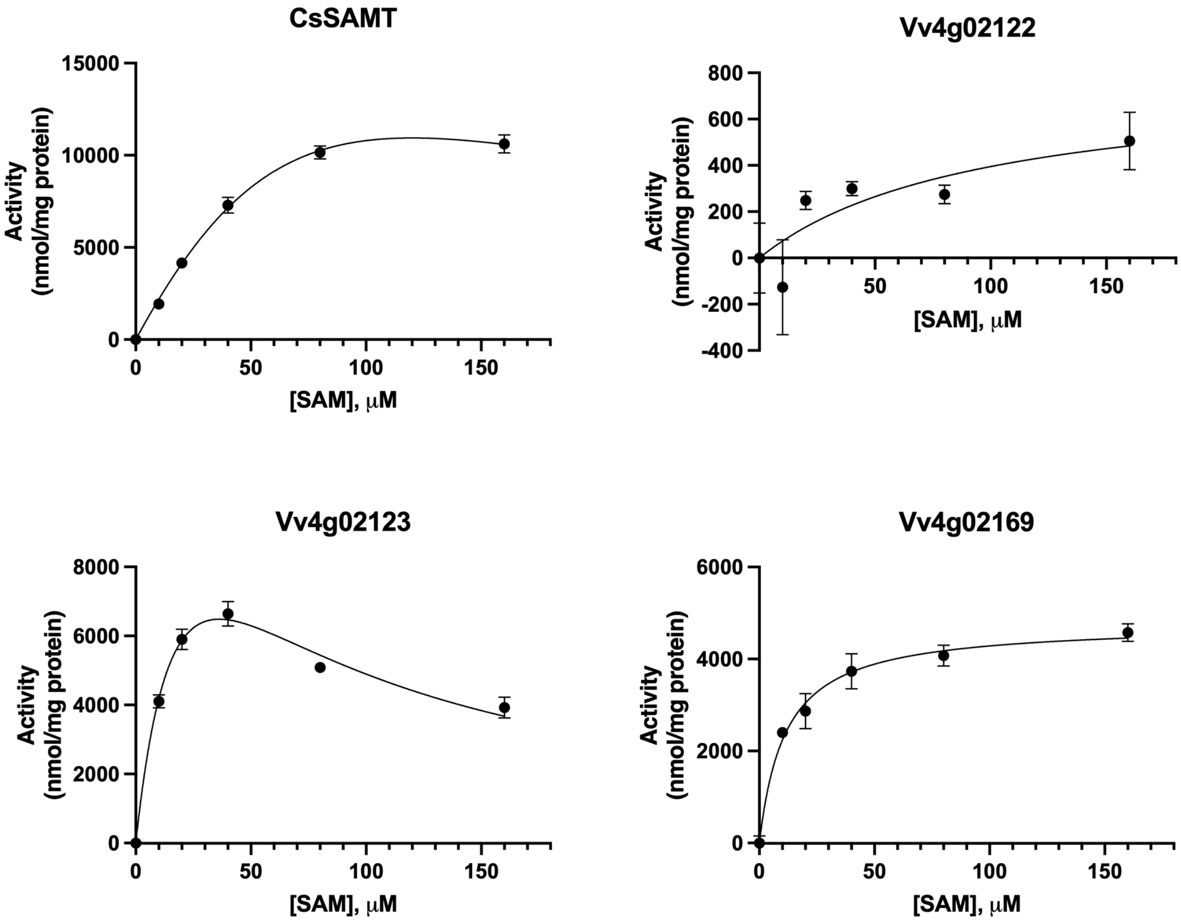


**Figure S16.** SAM kinetics plots of CsSAMT with saturating salicylic acid and Vv4g02122, Vv4g02123 and Vv4g02169 with saturating anthranilate. Error bars represent standard deviations; where error bars are not visible, standard deviation is too small to visualize. For all points, n=3.

**Table S1**. List of primers used for site-directed mutagenesis.

| | Primer | Sequence | Annealing Temperature (℃) |
|---|---|---|---|
| ZmAAMT1 | F164S-F | 5'-TGGCTACGCCATTGTAAACAACTCAACGAATGAAACAGGTGTAC-3' | 55 |
| | F164S-R | 5'-GTACACCTGTTTCATTCGTTGAGTTGTTTACAATGGCGTAGCCA-3' | |
| | F164Y-F | 5’-CACCTGTTTCATTCGTTGTATTGTTTACAATGGCGTAGC-3’ | 55 |
| | F164Y-R | 5’-GCTACGCCATTGTAAACAATACAACGAATGAAACAGGTG-3’ | |
| | C165S-F | 5’-GTTTCATTCGTTGTTTAGCTTACAATGGCGTAGC-3’ | 57 |
| | C165S-R | 5’-GCTACGCCATTGTAAGCTAAACAACGAATGAAAC-3’ | |
| | Q167W-F | 5’-CGTTGTTTTGTTTATGGTGGCGTAGCCAAG-3’ | 55 |
| | Q167W-R | 5’-CTTGGCTACGCCACCATAAACAAAACAACG-3’ | |
| | Y246H-F | 5'-GGGCCAACAGGCCATGCAGGTGATTGCTTTC-3' | 55 |
| | Y246H-R | 5'-GAAAGCAATCACCTGCATGGCCTGTTGGCCC-3' | |
| | Y246W-F | 5’-GAAAGCAATCACCTGTGGGGCCTGTTGGCCCAAAG-3’ | 55 |
| | Y246W-R | 5’-CTTTGGGCCAACAGGCCCCACAGGTGATTGCTTTC-3’ | |
| | I274Q-F | 5'-CGCCGACGCTGGGGCTATACTGTGGCAGGTAAAAGCTTTCC-3' | 63 |
| | I274Q-R | 5'-GGAAAGCTTTTACCTGCCACAGTATAGCCCCAGCGTCGGCG -3' | |
| | L329M-F | 5'-ATCACGGCACGCATACATTTGGCCACGTTCTCGCC-3' | 55 |
| | L329M-R | 5'-GGCGAGAACGTGGCCAAATGTATGCGTGCCGTGAT-3' | |
| | M333A-F | 5’-GTTTACGTGCCGTGGCGGAACCATTGGTAGC-3’ | 55 |
| | M333A-R | 5’-GCTACCAATGGTTCCGCCACGGCACGTAAAC-3’ | |
| | M333T-F | 5'-GCTACCAATGGTTCCGTCACGGCACGTAAACATTT-3' | 55 |
| | M333T-R | 5'-AAATGTTTACGTGCCGTGACGGAACCATTGGTAGC-3' | |
| | H368F-F | 5’-CGAAAAGACGAAATTTGCCGTGCTGGTGC-3’ | 72 |
| | H368F-R | 5’-GCACCAGCACGGCAAATTTCGTCTTTTCG-3’ | |
| | H368T-F | 5'-CACCAGCACGGCGGTTTTCGTCTTTTCGTTTTCCAGATGC-3' | 55 |
| | H368T-R | 5'-GCATCTGGAAAACGAAAAGACGAAAACCGCCGTGCTGGTG-3' | |
| | H368Y-F | 5'-GCACCAGCACGGCATATTTCGTCTTTTCGTTTTCCAGATGC-3' | 55 |
| | H368Y-R | 5'-GCATCTGGAAAACGAAAAGACGAAATATGCCGTGCTGGTGC-3' | |
| CsSAMT | Q159H-F | 5'-GAACTTGGCTTAACCAATGCAGGGAATATGAGCTATGAA-3' | 59 |
| | Q159H-R | 5'-TTCATAGCTCATATTCCCTGCATTGGTTAAGCCAAGTTC-3' | |
| | Q263C-F | 5'-TTTCTGCAGGACTTGGCGTATAGCATGGAATATTAAAGCAATTGACTTTCTCTTCC-3' | 61 |
| | Q263C-R | 5'-GGAAGAGAAAGTCAATTGCTTTAATATTCCATGCTATACGCCAAGTCCTGCAGAAA-3' | |
| | C319S-F | 5'-TGCTACCGCACGCATACTATTCGCAACATTATAGC-3' | 57 |
| | C319S-R | 5'-GCTATAATGTTGCGAATAGTATGCGTGCGGTAGCA-3' | |
| | M320I-F | 5'-TGCTACCGCACGTATACAATTCGCAACATTATAGCC-3' | 59 |
| | M320I-R | 5'-GGCTATAATGTTGCGAATTGTATACGTGCGGTAGCA-3' | |
| | M320L-F | 5'-CTACCGCACGCAAACAATTCGCAACATTATAGCCGC-3' | 67 |
| | M320L-R | 5'-GCGGCTATAATGTTGCGAATTGTTTGCGTGCGGTAG-3' | |
| | A324M-F | 5'-GACACCAGTAAGGGCTCCATTACCGCACGCATACAATTCGC-3' | 61 |
| | A324M-R | 5'-GCGAATTGTATGCGTGCGGTAATGGAGCCCTTACTGGTGTC-3' | |
| | A324T-F | 5'-CCAGTAAGGGCTCGGTTACCGCACGCATAC-3' | 62 |
| | A324T-R | 5'-CCAGTAAGGGCTCGGTTACCGCACGCATAC-3' | |
| | F359H-F | 5'-GTCAGAGAAACAGTGACGTTAATATGTTTAGTTTTCTCCTTGCTCATACG-3' | 59 |
| | F359H-R | 5'-CGTATGAGCAAGGAGAAAACTAAACATATTAACGTCACTGTTTCTCTGAC-3' | |
| | M320L/A324M-F | 5'-GACACCAGTAAGGGCTCCATTACCGCACGCAAACAATTCGC-3' | 64 |
| | M320L/A324M-R | 5'-GCGAATTGTTTGCGTGCGGTAATGGAGCCCTTACTGGTGTC-3' | |
| Vv4g02169 | T177S-F | 5'-GAACAAACAAATTTCGAACAAGGGG-3' | 50 |
| | T177S-R | 5'-CCCCTTGTTCGAAATTTGTTTGTTC-3' | |
| | G281E-F | 5'-GAGGATTTGGAGACAGAAATCGAGAATGATGG-3' | 55 |
| | G281E-R | 5'-CCATCATTCTCGATTTCTGTCTCCAAATCCTC-3' | |
| | E358D-F | 5'-CACTAGAGAAGTGGATCACGTGAGTGTTCTG-3' | 56 |
| | E358D-R | 5'-CAGAACACTCACGTGATCCACTTCTCTAGTG-3' | |

**Data S1.** Codon-optimized gene sequences of AAMTs and SAMTs.

>CsSAMT

ATGGAAGTGGTGCAGGTCTTACACATGAACGGCGGAGTTGGTAATGCTTCGTACGCAAGTAATTCGCTG
GTGCAAAAGAAGGTTATTTCCATTGCTAAGCCTATAACCGAAGAGGCCATGACCAAATTATTTTGCAGTA
CCTTTCCTACTAAAGTAGCAATAGCAGATCTGGGTTGTTCTAGTGGTCCGAATACATTGCTCGTCGCGTC
TGAGTTAATCAAGGTTGTAAATAAAATATGTGATAAACTGGGGTCTCAGCTGCCGGAATTTCAGGTTTTC
CTGAATGACCTGCCCGGAAACGATTTTAATACGATCTTTCGTTCACTGGCCAGCTTCCAAAAGATTCTTC
GAAAACAACTTGGCTCAGCATCAGGTGCAGCGGGCCAATGTTTCTTCACAGGCGTTCCGGGGTCATTT
TACGGCAGATTATTCCCGCGCAACTCCGTTCACCTCTTTCATAGCTCATATTCCCTGCAATGGTTAAGCC
AAGTTCCGGATGGCTTAGAGTCTAACAAAGGAAATATCTTTATGGCATCCACCAGCCCGCCTTGTGTCC
TGACCGCTTACTACGAACAGTTTCAAAGAGACTTTTCACTCTTCTTGAAATGCAGATCAGAGGAACTGG
TGGCTGAGGGAAGAATGGTACTTACATTTCTGGGCCGGAAGAGCCAGGATCCTTCATCGAAGGAATGC
TGCTATATATGGGAGTTACTGGCAACGGCGTTAAATAATATGGTCAGTGAAGGTCTCATCGAGGAAGAGA
AAGTCAATTGCTTTAATATTCCACAATATACGCCAAGTCCTGCAGAAATCAAGTCGGAAGTAATTAAGGAA
GGAAGCTTCACGATTGATCACCTGGAAGTTAGCGAAGTCAATTGGAACGCCTACCAGAACGGTTTTAAG
TTCAATGAGGCCGTTGATGCGTTTAACGACGGCGGCTATAATGTTGCGAATTGTATGCGTGCGGTAGCA
GAGCCCTTACTGGTGTCTCAGTTTGGCGAGAGTATCATCGACGAGCTTTTCAAACGCTATCGGGAGATA
GTTGCGGATCGTATGAGCAAGGAGAAAACTAAATTTATTAACGTCACTGTTTCTCTGACCAAGATAGGGT
GA

>Vv4g02122

ATGGAAGTGGTACAAGTACTCTGCATGAAGGGTGGCAATGGAGATACTAGTTATGCGCAAAACTCCCTT
TTACAAAAGAAGGTGATCAGCCTCACAAAGCCTATCACCGACGAAGCTATTTCTAATCTTTTCTGCAATA
ACTTCCCAGCCCGTCTGTGCATAGCTGACCTTGGATGCAGCTCCGGGCCGAATACGTTGTTTGCTGTT
CTGGAGTTCGTCACGACGGTGGACAAGGTCCATAAGAAGATGGGGCACGAGCTCCCTGAGATCCAAG
TCTTTCTTAACGACTTACCAGGAAATGACTTTAACACGATCTTCAAGTCACTTCCAACGTTCCAAAAGGA
TTTACAAAAGACCATGGGTGCGGGCGCCGAATCTTGCTTCGTTACTGGTGTACCGGGCAGTTTCTACG
GTCGCTTGTTCCTTGGCAAGTCCTTGCATTTTGTTCACAGCTCATACAGTCTCCAGTGGCTTTCACAAG
TTCCGCGAGGCTTGGAATCGAACAAGGGGAACATTTACATGGCATCATCGTCCCCACCGAGTGTTCTG
AAGGCTTACTATGAGCAGTTCCAAACGGACTTCTCAATGTTCCTGCGATGTCGGAGTGAGGAGCTGTT
GGAAGGTGGCAGTATGGTTCTCACATTCCTGGGGCGCATTTCTGAAGATCCGTCCAGTAAAGAATGCT
GCTACATCTGGGAACTGCTTGCAGTTGCTCTCAACGACATGGTGGCAGAGGGGCTTGTCGAGGAAGA
GAAAATGGACAGTTTCAACATACCTCAATATACTCCTTCACCGGCTGAGGTCAAGTGTGAGGTGGAGAA
AGAGGGGAGCTTCACAATCAATCGATTAGAGGCTTCTGAAGTGAACTGGAATGCATATCACGGTGAGTT
TTGTCCTTCCGATGCGCACGAGGACGGCGGATACAATGTAGCGAAGCTGATGCGTGCGGTCGCCGAG
CCGCTTCTGGTGAGCTACTTCGGCGACGGCATAATTGAAGAGGTTTTCTCTCGATACCAGAAGATTGTG
GCCGACCGGATGAGTCGTGAGAAGACTGAGTTTGTGAATGTAACGGTAAGTATGACTAAGCGCGGATG
A

>Vv4g02123

ATGGAAGTTGTACAGGTTCTCTGCATGAAAGGTGGAAACGGTGATACGTCTTACGCTAAGAACTCTCTG
GTACAAAAGAAGGTCATCTCTTTGACCAAACCGATTATTGAAGAAGCCATAACTAACCTTTATTGCAATAA
ATTCCCTACCTCTCTCTGCATCGCGGACCTGGGCTGCAGTAGTGGTCCTAACACGCTTTTCGCAGTTCT
TGAGGTAGTCACTACCGTGGACCGTGTTGGCAAGAAGATGGGACGTCAACTTCCAGAGATTCAGGTCT
TTCTGAATGATCTCCCCGGCAACGACTTCAATACTATATTTAAAAGTTTACCCCGTTTCCAAAAGGACTTG
GAGAAGCGCATGGGTGCGGGCGCGGAGAGCTGTTTCATAAACGGCGTTCCCGGTTCCTTTTACGGCC
GGTTATTCCCTTCTAAGTCATTGCATTTCATACATAGTTCCTACAGCCTGCAGTGGCTGTCACAAGTTCC
CCAAGGCTTAGAGTCGAATAAGGGCAACATTTATATGGCCAGCTCTTCTCCGCCCTGCGTATTAAAAGTC

TACTATGAGCAATTCCGTACCGACTTCAGCATGTTCCTGCGTTGTCGCTCAGAGGAGTTACTGGAAGGC
GGTTCAATGGTCTTGACATTCTTGGGTCGTCGTAGCGAGGACCCCTCATCAAAGGAGTGCTGTTACATC
TGGGAACTTTTGGCTGTAGCCTTAAATGATATGGTTGCAGAGGGATTAATAGAGGAAGAGAAGATGGATT
CCTTCAACATCCCGCAATACACTCCGAGCCCGGCTGAGGTGAAGTGTGAGGTGGAGAAGGAAGGAAG
CTTCACCATCTCAAAACTTGAGGTTTCGGAAGTCAATTGGAACGCTTACCATGGTGAGTTCTGCCCTTC
TGATGCGCACAAGGATGGTGGTTATAACGTGGCGAAGTTAATGCGGGCTGTGGCGGAGCCGTTACTGG
TTTCACACTTCGGAGATGGAATCATTGAGGAAGTCTTTTCTCGATACCAGAAGATAGTTGCAGACCGCAT
GTCCCGCGAAAAGACCGAATTCGTTAATGTCACTGTGTCCATGACGAAGCGTGGCTGA

>Vv4g02169

ATGAAGAAGAAAGAGAGCATGGGTGTCCAACAGGTAATTTGCATGAAAGGCGGCGTGGGAGAAGGAA
GCTATGCTCGTAACTCTAAGTCCCAAGCGGCGCTCTTGTCGAAGTCTATGCCCTTGTTAGAGCAAGCGG
TCCTGGACCTCTGTTGTACCACCCTTCCAGAGAGCGTGGCAATAGCAGACCTGGGTTGTTCCTCGGGA
CCAAATACATTCTTCGCCGTCTCAGAGATCATGACCATAATATACCGTCGTTGTCGCCAACTGGGCCGCT
CGCCGCCGGGCTTCTGGGTCTTCCTGAACGACTTGCCAGGAAATGACTTCAACGCAGTGTTCAAGTC
GCTCCCAACATTCCACGAGAAGATGAAGGAAGAGAACGGACAGGAGTTCGGACCTTGTCATGTGGCG
GCGGTGCCGGGTAGCTTCTACCACAAGCTGTTCCCTTCTCGACGCTTGCATTTCGTCCATAGTTCTTGC
TCCTTGCACTGGCTGTCCCAAGTTCCACCAGAATTACTGAACAAACAAATTACGAACAAGGGGAAAATT
TATCTCTCTAAAACCTCAAGTCCCGCCCTCATAGACGCATACGCGAGCCAATTCCAGCGAGACTTCAGT
CTGTTCTTGAAGTTGCGTTCAGAGGAGACTGTGCCAGGCGGCAGAATGGTTCTGAGCTTGATGGCAC
GTCGTACGCCTGATCCAGTATCCGATGAGAGTTGTCTCCTTTGGGATCTCTTGGCCCAGGCGCTGCAA
GGCTTAGTTTCTGAAGGGCTCATTGCGGAAGAGAAATTGGATTCGTACAATGCTCCATACTACCAGCCA
TACACTGAGGATTTGGAGACAGGAATCGAGAATGATGGTAGTTTTAGTATCAACGGTTTAGAGATAATGG
TATTGCCGTGGGACTCCGCCTCAGGTGGTCAAAACTACGATCGCCCAACGACCGCTCAAAAGATCGCG
AAGTCCATGAAGGCCGTCCAAGAGCCAATGCTGGCGTCCCACTTCGGAGCGGAAATCATGGACCCAC
TGTTTAAGAGACTGATGGAAATTATAGCCGCAGACACTAGAGAAGTGGAACACGTGAGTGTTCTGGTGA
GCATGACTCGTAAAGCTTAA

>Vv12g00725

ATGGAAGTCCAACAGGTCCTGTGTATGAAGGGTGGTGATGGTGAGGCTTCGTACGCAAATAACAGTCTT
CTGCAAAAGAAAGTAATATTGGAAGTAAAGCCGATCTTGGAAGAGTCTATCACGGAATTATACTGCAAGA
CCTTTAGCGAGTGCTTAAAGATAGCGGACCTTGGATGCTCTTCCGGCCCCAACACCTTCTTACCCTTAT
GGGAGATTATTGATTGCATCGGAGCAACCTGTAGTCGTTTTAGCCGTGAGCCGCCTGCCTTCCAGATTT
TCCTTAACGACTTACCACAAAATGACTTCAACGCAATTTTCGAGAGCCTGGCTAGATTCTATGAGCGCAT
CGAGAAGGAAAAGGAAGGGATGTCACGTCAATGCTTTATAGCGGGAGTTCCTGGTTCATTCCATCGTAG
ACTGTTTCCAGACCGATCTATTCACTTCTTTCACTCATCGTACTCTCTGCACTGGCTTAGCCAGGTACCG
GAAGGGCTTGTATCTGAATCGGGCACTCCTTTGAATAAGGGTAACATTCACCTGACAGTTACTACTCCA
CCCAGTGTTCACAAGGCATACTTAAACCAATTCGAACGCGACTTCACAGCATTCTTGAGATTACGTTCTC
AGGAAATAATACCTGGCGGGCATATGTTACTGACGCTCTTAGGTTCCGATGGCAATGGTCAGAACTCTT
CCACCGATGGGTTATACAAGATCTGTGAATTGATAAGCATGACACTGAAGGATATGGTAACTGAAGGGTC
AATACAAGAATCTGAGCTGGACTCTTTAAACATTCCTTTATTCATGCCTTCTCCGGAGCAAGTTAGAAGT
GTCATTCAGCGCGAGAGCTCATTCACCTTATTGCGTCTTGAAACTTTCAAGTTAGATTGGGCTGACAAC
ATCGATGACGGTAACAAAGACCAAGTTTTCGATAAGTACGGCAGAGCGAAGTACGTCGTTATGTACATC
CGTGCAGTCGGCGAACCCATCCTGGCCTCACATTTCGGCGGTGCCGTTATGGACTCCTTGTTCCACAG
ATTCTTTATGAAGGTAGTAGAAAATATTGAGACGGGAAAGGGTATCTACACGAACCTCGTCATTAGTCTG
TCACGGAACGGTAGCCTGCCAACCATGGATCGTAAGGGGTAA

>Concord_4g02123

ATGGAAGTGGTACAAGTGCTGTGCATGAAGGGTGGCAACGGAGATACTTCCTACGCCAAGAATAGTTTA
GTCCAAAAGAAGGTGATCAGCCTTACTAAGCCTATTATCGAGGAAGCTATAACGAATCTTTACTACAACA
AATTCCTCACTAGTTTGTGCATAGCTGACCTTGGGTGCTCATCAGGCCCGAACACACTTTTCGCGGTAT
TGGAAGTTGTTACCACCGTAGATCGTGTGGGAAAGAAGATGGGCAGACAACTTCCCGAGATACAAGTC
TTCTTGAATGACCTGCCGGGGAATGACTTTAACACTATCTTTAAGTCACTCCCAGGCTTCCAGAAAGAC
TTAGAAAAGCGCATGGGTGCAGGCGCCGAGTCATGCTTCATTAACGGCGTCCCCGGATCGTTCTATGG
GAGACTCTTCCCATCAAAGTCCCTGCACTTCATTCACAGCAGCTACTCTCTTCAATGGCTTTCGCAGGT
TCCCCAGGGTCTTGAGAGCAATAAGGGAAATATTTATATGGCAAGCAGTAGCCCACCGTGCGTACTGAA
GGTTTACTACGAACAATTCCGTACTGATTTCTCCATGTTCTTGCGTTGTCGTTCAGAAGAGCTTTTGGAA
GGCGGTAGTATGGTATTAACTTTTCTTGGGCGGCGTTCAGAGGACCCAAGTTCGAAGGAATGTTGCTAT
ATATGGGAGCTCTTAGCCGTTGCTCTGAACGACATGGTTGCGGAAGGTTTAATCGAAGAGGAGAAGAT
GGACAGTTTCAATATCCCACAATACACACCTTCTCCGGCTGAGGTGAAGTGTGAGGTAGAGAAGGAAG
GAAGCTTCACCATCTCACGCTTAGAGGTAAGTGAGGTTAACTGGAACGCGTACCATGGGGAGTTCTGT
CCGAGTGACGCACATAAAGATGGTGGGTACAATGTCGCGAAGTTAATGCGAGCAGTAGCCGAACCACT
TCTCGTCTCGCACTTCGGTGACGGCATCATCGAGGAAGTATTCTCACGTTACCAAAAGATTGTTGCTGA
CCGCATGAGCAGAGAGAAGACCGAGTTTGTGAATGTCACCGTTAGCATGACCAAACGCGGTTGA

>ZmAAMT1

ATGCCCATGCGCATTGAACGCGACTTACACATGGCTATTGGTAATGGCGAGACCTCTTATACCAAGAAC
AGTCGCATCCAGGAAAAGGCGATGTTCCAAATGAAAAGCGTTCTGGAAGAAGCGACCCGCGCCGTCT
GTACCACCCTTTTGCCGCAGACAATGGTGGTAGCAGATCTGGGTTGTAGCTCCGGCCCGAATACCCTT
CGTTTTGTAACGGAAGTTACCCGCATTATTGCGCATCACTGTAAACTTGAACATAATCGCCGTCACGATC
ATTTGCCTCAACTGCAATTCTTCCTCAACGATTTGCCTGGCAATGATTTTAATAACCTGTTTCAACTGATT
GAACAATTTAACAAATCTTCAACGACCCATAAAGGCGACGCCGCGACGGAAGCGCTGCAACCGCCGTG
TTACATTTCAGGCTTACCAGGAAGTTATTATACGCGCATCTTTAGCTCGGAGTCGGTACACCTGTTTCATT
CGTTGTTTTGTTTACAATGGCGTAGCCAAGCGCCGGAGCAGTTAAAAGGTACACAGAAGAGCTGTCTG
GACATTTATATTACGAAAGCGATGTCGCCGTCCATGGTAAAACTTTTCCAGCAACAATTCCAGAAAGATT
TTAGCCTTTTCTTGCGTCTTCGTTACGAAGAGCTGGTCTCGGGTGGTCAGATGGTATTAACCTTCATCG
GCCGTAAACACGAAGACGTTTTCACGGGCGAAAGCAATCACCTGTATGGCCTGTTGGCCCAAAGTTTA
AAGTCATTGGTGGACGAAGGGCTGGTTGAAAAGGAGAAGCTGGAAAGCTTTTACCTGCCAATTTATAGC
CCCAGCGTCGGCGAGGTTGAAGCAATCGTCAAACAGCTCGGTCTGTTTAATATGAACCACGTGAAGGT
TTTCGAAATTAACTGGGACCCGTATGACGATAGTGAGGGCGACGACGTACACAATTCTATCGAATCAGG
CGAGAACGTGGCCAAATGTTTACGTGCCGTGATGGAACCATTGGTAGCCTCGCAGTTCGGCGAGCGTA
TTCTGGATGAATTGTTTAAGGAATATGCCCGTCGCGTCGCGAAGCATCTGGAAAACGAAAAGACGAAAC
ACGCCGTGCTGGTGCTGTCGATTGAAAAGGCGATTATCCACGTCTAA
